# Joint phylogenetic estimation of geographic movements and biome shifts during the global diversification of *Viburnum*

**DOI:** 10.1101/811067

**Authors:** Michael J. Landis, Deren A. R. Eaton, Wendy L. Clement, Brian Park, Elizabeth L. Spriggs, Patrick W. Sweeney, Erika J. Edwards, Michael J. Donoghue

## Abstract

Phylogeny, fossils, biogeography, and biome occupancy provide evidence that reflects the singular evolutionary history of a clade. Despite the connections that bind them together, these lines of evidence are most often studied separately, by first inferring a fossil-dated molecular phylogeny, then mapping on ancestral ranges and biomes inferred from extant species. Here we jointly model the evolution of biogeographic ranges, biome affinities, and molecular sequences, incorporating fossils to estimate a dated phylogeny for all of the 163 extant species of the woody plant clade *Viburnum* (Adoxaceae) that we currently recognize. Our analyses indicate that while the major *Viburnum* lineages evolved in the Eocene, the majority of extant species originated since the Miocene. *Viburnum* radiated first in Asia, in warm, broad-leaved evergreen (lucidophyllous) forests. Within Asia we infer several early shifts into more tropical forests, and multiple shifts into forests that experience prolonged freezing. From Asia we infer two early movements into the New World. These two lineages probably first occupied warm temperate forests and adapted later to spreading cold climates. One of these lineages (*Porphyrotinus*) occupied cloud forests and moved south through the mountains of the Neotropics. Several other movements into North America took place more recently, facilitated by prior adaptations to freezing in the Old World. We also infer four disjunctions between Asia and Europe: the *Tinus* lineage is the oldest and probably occupied warm forests when it spread, while the other three were more recent and in cold-adapted lineages. These results variously contradict published accounts, especially the view that *Viburnum* radiated initially in cold forests and, accordingly, maintained vessel elements with scalariform perforations. We explored how the location and biome assignments of fossils affected our inference of ancestral areas and biome states. Our results are sensitive to, but not entirely dependent upon, the inclusion of fossil biome data. We argue that it will be critical to take advantage of all available lines of evidence to decipher events in the distant past, and the joint estimation approach developed here provides cautious hope even when fossil evidence is limited.

## INTRODUCTION

Convincing historical explanations for the modern distribution of organisms are hard to come by. This is because they necessitate, at least: (1) a compelling phylogenetic analysis of a comprehensive sample of relevant species (a phylogenetic component); (2) the incorporation of fossils to assess the absolute timing of events (a temporal component); (3) a historical biogeographic analysis that identifies and explains geographic disjunctions (a spatial component); and (4) information on the environments occupied by the species under consideration (an ecological component). Using quantitative phylogenetic models, numerous recent studies have extracted important insights from the comparison of dated phylogenies, geographic inferences, and biome reconstructions (e.g., Weeks et al., 2014; Meseguer et al., 2015; Cardillo et al., 2017; Gagnon et al., 2019). On the whole, analyses of these components have been carried out sequentially: species relationships are first inferred from sequence data, the tree is then time-calibrated with fossils, and the dated tree is used to separately infer geographic movements and biome shifts during the evolution of a clade. With sequential analyses, by design, evolutionary inferences that are made during the later stages cannot influence inferences earlier in the analytical sequence.

It is possible, of course, that in practice, molecular and morphological evidence will often be sufficient to infer phylogenetic relationships or divergence times during that first stage of analysis. But perhaps not always. For example, the movement and diversification of lineages are influenced by the expansion and contraction of biomes. And, from an evidential standpoint, a fossil may have too few morphological characters to securely position it during the first (phylogenetic) stage of analysis. Yet, phylogenetically unresolved fossils still indicate that at least one ancestral lineage was associated with a particular region or biome, even if it is uncertain exactly which lineage that is. Furthermore, the temporal, geographic, and ecological evidence that fossils provide may also aid in their phylogenetic placement.

Here we implement an approach that jointly models the evolution of molecular sequences, fossils, geographic ranges, and biome affinities, and in which these collectively determine an outcome. Conceptually, we view this approach — the merging of these key elements into a single analytical framework — as an extension to biogeography of the “combined evidence” paradigm developed for “tip-dating” phylogenies (Nylander et al., 2004; Pyron, 2011; Ronquist et al., 2012). The combined evidence strategy favors evolutionary histories in harmony across phylogenetic, temporal, biogeographic, and ecological components, while penalizing those with dissonant components.

The specific question we address here is how *Viburnum* (Adoxaceae, Dipscales) – a widespread angiosperm lineage of ~163 species of shrubs and small trees – radiated into the geographic ranges and biomes that it occupies today. We and others have published on this problem previously, so our analyses provide a critical test of recently proposed and strongly contrasting hypotheses. Furthermore, as *Viburnum* exemplifies Northern Hemisphere disjunctions that are common in plants, fungi, and animals, as well as transitions between the major mesic forest biomes (tropical, warm temperate, and cold temperate), our results address questions of long-standing interest to evolutionary biologists, biogeographers, and ecologists.

Apart from the details of this case, however, we hope that the approach featured here will illuminate an important point. It is, it must be admitted, a humbling task to infer ancient events, and the results in many cases are tenuous at best. Given the obvious limitations – working, on the whole, with extant species and few, if any, fossils– it is necessary to integrate all of the available sources of evidence if we hope to produce assuring answers. The consilience of multiple lines of evidence may give us a chance to convincingly favor some hypotheses over others, even in cases that appear unpromising on the surface.

### Background Information

*Viburnum* is well suited for such inference problems. First, we have tried to sample its species comprehensively, making many recent field collections throughout its wide geographic range and ensuring accurate identifications. Second, in addition to several major sources of molecular data, including whole chloroplast sequences (cpDNA; Clement et al. 2014) and nuclear genomic data (RAD-seq; Eaton et al. 2017), we have assembled extensive morphological information (Schmerler et al. 2012; Spriggs et al. 2018), including data on relevant functional traits (Chatelet et al. 2013; Edwards et al. 2014; Scoffoni et al. 2016). We have also carried out long-term observations in the Arnold Arboretum of Harvard University and at several field sites (Edwards et al., 2017), phylogeographic analyses within several species complexes (Spriggs et al. 2019a,b; Park and Donoghue 2019), and analyses of relevant climate variables at several scales (Edwards et al. 2017; Spriggs et al., 2019; Park and Donoghue, in review). Third, specific hypotheses have recently been advanced about the biogeographic and ecological history of *Viburnum* (beginning with Winkworth and Donoghue 2005; Moore and Donoghue, 2007, 2009; Clement and Donoghue 2011, Clement et al. 2014). On the one hand, it has been argued that *Viburnum* originated in tropical forests, and that the few species present today in these forests are “dying embers” of a deep tropical past (Spriggs et al. 2015). On the other hand, it has been hypothesized that *Viburnum* originated in cold forests that experience prolonged freezing, and that they spread only later into tropical environments (Lens et al. 2016). These starkly contrasting views have a direct bearing on how we interpret the evolution of morphological and functional traits, especially the directionality of leaf and wood evolution.

As shown in Figure 1, Eastern Asia is the center of extant *Viburnum* species *and* phylogenetic diversity, with roughly 89 species representing 12 of the major clades marked in Figs. 3, 4, and 5. These Asian species extend from nearly aseasonal tropical forests (sometimes at low elevations in Indonesia and Malaysia), northward through monsoonal broadleaved evergreen (lucidophyllous) forests in India, Vietnam, Taiwan, and southern China, and into the colder temperate forests of China, Japan, and Korea, and the boreal forests of Northern Japan and Russia. *Viburnum* also extends westward across the Himalayas, with one species (*V. cotinifolium*) reaching Afghanistan.

**Figure 1.**
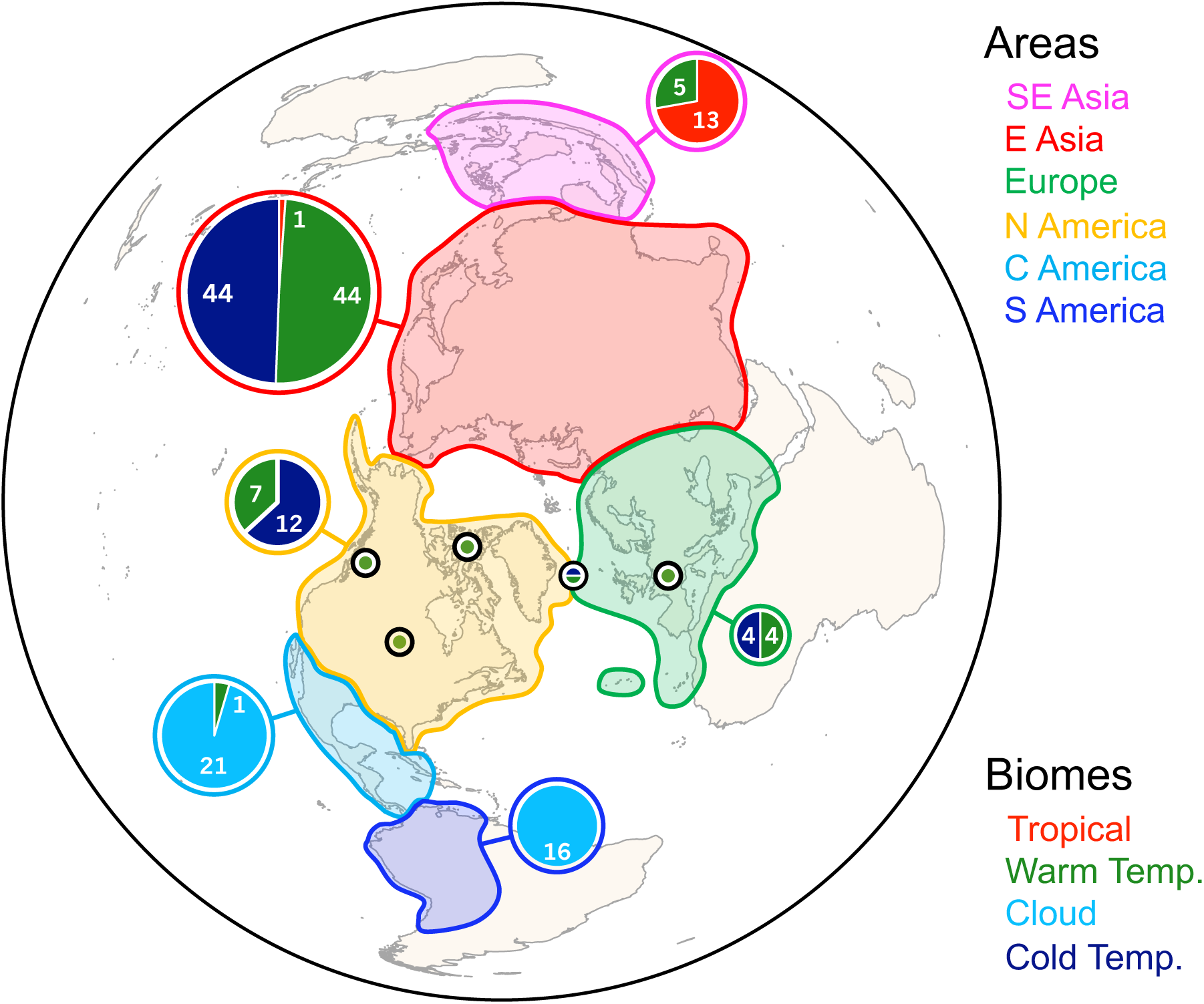
Global diversity of *Viburnum* across six areas and four biomes. Areas are marked with colored polygons for Southeast Asia (magenta), East Asia (red), Europe (green), North America (yellow), Central America and Mexico (cyan), and South America (blue). Counts of local species with biome affinities are reported for each area with pie charts, with the area of each chart corresponding to total number of local species. Biomes are colored as tropical (red), warm temperate/lucidophyllous (green), cloud forest (sky blue), and cold temperate forest (dark blue). Asterisks represent the five fossil pollen localities.

In Europe there are eight extant species belonging to four major clades (Clement et al., 2014). *Viburnum tinus* is widespread around the Mediterranean (including northern Africa), while three other species extend into temperate (*V. lantana*, *V. maculatum*) and boreal (*V. opulus*) forests. Narrowly endemic species are found in the Caucasus Mountains (*V. orientale*), the Canary Islands (*V. rugosum*), and the Azores (*V. treleasei*).

In the New World north of Mexico, there are 19 species in five major clades. Only one species, *V. ellipticum*, is endemic to the northwestern USA. In eastern North America, viburnums extend from subtropical forests in Florida through temperate and boreal forests as far north as Newfoundland. A clade of ~36 species occupies cloud forests in the mountains of Mexico and Central America, the Andes of South America (south to northern Argentina), and the Caribbean islands of Jamaica and Cuba.

Similar biogeographic and ecological patterns are common across angiosperms. As is the case for many Laurasian clades in mesic environments, we also note that there are major regions and biomes that are unoccupied by *Viburnum*. In particular, no species are native to sub-Saharan Africa, Madagascar, Australia, or New Zealand, and they are absent from deserts, grasslands, and other arid biomes. Likewise, the traits that vary extensively in *Viburnum* are of general interest to botanists, including leaf form, wood anatomy, and leafing habit (deciduous versus evergreen). However, for the record, other characters vary little in *Viburnum*: they have small white flowers pollinated mainly by bees and flies, and fleshy single-seeded fruits, dispersed mainly by birds. None are herbaceous, succulent, or spiny plants, and few reach heights of over 20 meters.

## MATERIALS AND METHODS

### Analysis overview

Our goal was to estimate a joint distribution of phylogenetic topologies, time-calibrated lineage splitting events, biogeographic histories, and biome shift histories for 163 extant *Viburnum* species and 5 fossil specimens. To this end we adopted a two-stage approach. During Stage 1, we estimated a tree topology for *Viburnum* from RAD-seq data for 118 species using a phylogenomic concatenation method. In Stage 2, we estimated absolute divergence times, ancestral ranges, and ancestral biomes while requiring that all 163 extant species relationships be congruent with the 99%-majority rule consensus tree from Stage 1. To do so, we jointly modeled the evolution of the biogeographic ranges, the biome affinities, and ten partitioned molecular loci (9 cpDNA and nrITS) to estimate node ages and any equivocal species relationships. Dating information enters the model through five fossil occurrences, generated by the fossilized birth-death process, and through a secondary calibration for the origin of crown Adoxaceae. Morphology-based clade constraints are additionally imposed upon five fossil taxa and ten unsequenced extant species (Supplements 3, 4).

The second stage of our approach takes inspiration from morphology-based combined evidence (Nylander et al. 2004) and tip-dating (Pyron 2011, Ronquist et al. 2012) analyses, and from fossil-based ancestral state reconstructions for biogeography (Mao et al., 2012; Wood et al., 2012) and biome occupancy (Betancur et al., 2015). Both tip-dating and fossil-based ancestral state estimates incorporate fossil taxa as leaf nodes in the phylogeny. Rather than first estimating the phylogeny of extant and fossil taxa, and then estimating ancestral ranges and biome affinities, we directly include biome and biogeographic data as part of the combined evidence tip-dating exercise. This means that each of our joint posterior samples represents an internally consistent evolutionary scenario that could have unfolded under our generative model to produce the evidence we have collected.

### Stage 1: RAD-seq topology estimation

We estimated the *Viburnum* tree topology using newly sequenced RAD data for this study add to data from a previous study (Eaton et al., 2017), nearly doubling the number of taxa with genomic data from 64 to 127, including 9 outgroup species (Supplement 1). These newly sequenced species filled sampling gaps for many sub-clades; in addition, we added representatives of one major clade (*Mollotinus*: *V. ellipticum*, *V. molle*) and increased the sampling of the Neotropical *Oreinotinus* clade from 4 to 25 species. As in the previous study, RAD-seq libraries were prepared by Floragenex, Inc. (USA) using the PstI enzyme and size-selection for a mean fragment length of 400bp following sonication. The samples from this study are drawn from six different libraries. We found high consistency in the recovery of shared RAD loci across repeated library preparations, showing no evidence of plate-effects (e.g., increased missing data between samples prepared in different libraries). Raw RAD-seq data have been submitted to NCBI SRA under project number PRJNAXXX.

The RAD-seq dataset was assembled in ipyrad v.0.7.13 (Eaton 2014). Raw reads from all sequencing lanes were demultiplexed separately in ipyrad and then merged into a single assembly object, from which samples were sub-selected in a new branch in which a single set of parameters was applied to all samples. This included trimming reads for low-quality scores (Q ≤ 30) and Illumina adapters (filter adapters = 2), clustering reads at an 85% similarity threshold, and calling consensus bases and alleles for clusters with a depth ≥6 (mindepthstatistical = 6).

The RAD-seq topology for the 127 *Viburnum* and outgroup species was inferred using RAxML (Stamatakis, 2014) to estimate the tree and bootstrap values under a concatenated GTR+Gamma model. Outgroup species were then pruned from the estimated tree to yield the backbone topology for 118 of ~163 (72%) *Viburnum* species in Fig. 2A. Nodes with equivocal bootstrap support (p < 0.99) were collapsed into polytomies, resulting in the RAD-seq backbone topology used in Stage 2.

**Figure 2.**
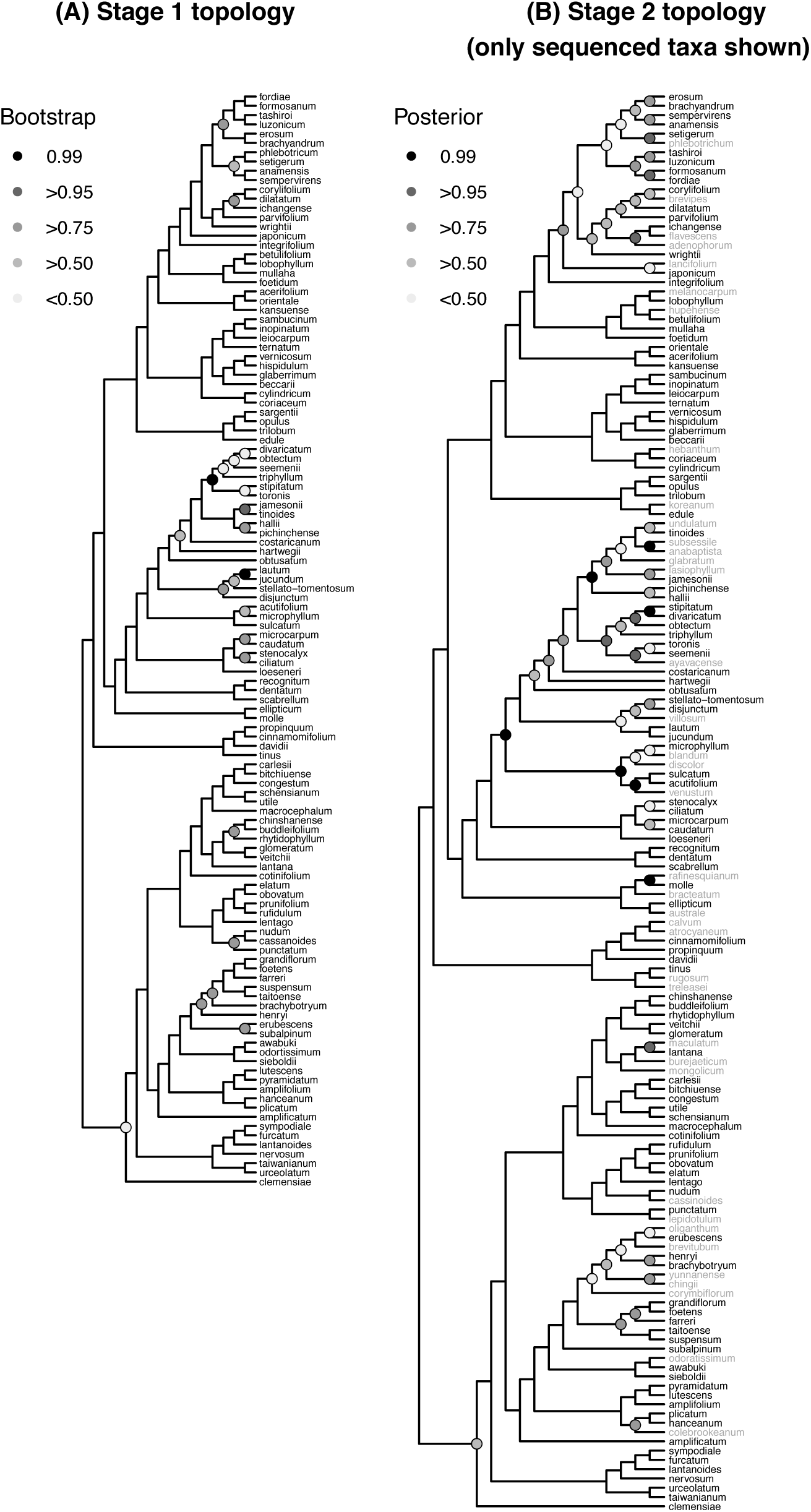
*Viburnum* topological estimates. (A) The Stage 1 topology represents the relationships of 118 species estimated from RAD-seq data using RAxML. Clades are annotated with bootstrap support. Clades with bootstrap support of 99 or 100 were then imposed as clade constraints for the Stage 2 analysis. (B) The Stage 2 topology represents the maximum clade credibility tree estimated using all molecular, fossil, geographical range, and biome occupancy data using the Bayesian framework described in the main text. Of the 168 taxa used in the Stage 2 analysis, 15 unsequenced (5 fossil and 10 extant) taxa were pruned from all posterior tree samples before constructing the consensus tree for the remaining 153 taxa. Clades are annotated with posterior probabilities (shaded markers). Taxon labels are colored black for taxa with RAD-seq and cpDNA data, and colored gray for taxa with only cpDNA data.

### Stage 2: Joint Bayesian macroevolutionary inference

We constructed a phylogenetic model to jointly estimate species relationships, divergence times, biogeographical histories, and biome shifts. Below, we first describe the data used for Stage 2, and then the components of the macroevolutionary model.

Molecular sequences for 153 of ~163 (94%) extant species were used to estimate species relationships among 35 lineages that were not topologically constrained and to estimate all phylogenetic divergence times under a relaxed clock model. We assembled previously published sequences for 145 extant viburnums for nine chloroplast genes (*mat*K, *ndh*F, *pet*BD, *psb*A, *rbc*L, *rpl*32, *trn*C, *trn*K, and *trn*SG) and one nuclear ribosomal marker (ITS). We then replaced 10 of those sequences that originated from herbarium or botanical garden specimens with new sequences extracted from field specimens that we collected. In addition, we sequenced 8 previously unsequenced species for this study, bringing the total number of species for these data to 153. The final matrix contained 22.6% missing cells. Sequences were aligned in Muscle then manually adjusted in AliView as needed. Sequence accession numbers are given in Supplement 2.

Geographical regions and biome affinities were scored for all 163 extant species and for the 5 fossil specimens. All species ranges were coded into one or more of six discrete areas: Southeast Asia, Eastern Asia, Europe, North America, Central America, and South America. These areas are meant to reflect the major centers of endemism for plant clades distributed around the Northern Hemisphere (Laurasian distributions), however we omitted Western North America as only one species of *Viburnum* (*V. ellipticum*) is endemic to that region. We subdivided Asia into Eastern and Southeastern regions to reflect patterns of endemism in *Viburnum*, especially the distribution of a number of lineages in Southeast Asia (Vietnam, Malaysia, Indonesia, and the Philippines). We subdivided the Neotropics into Central and South America both to reflect the isolation of South America before the Miocene and to assess the pattern and timing of the occupation of South America.

Species were assigned affinities to one or more of four mesic forest biome classes, following Edwards et al. (2017): tropical/subtropical (T), warm temperate (i.e., lucidophyllous) (W), cold temperate (Co), and/or cloud forest (Cl). Tropical forests experience limited seasonality (i.e., temperature and precipitation remain relatively stable and high throughout the year). Warm temperate forests are characterized by greater seasonality (warm/wet versus cool/dry seasons), but do not experience a period of prolonged freezing, and they are dominated by broad-leaved evergreen trees. In contrast, cold temperate forests experience a season in which temperatures routinely fall below freezing for one or more months, and are dominated by conifers and deciduous angiosperms. Cloud forests occur in montane regions at tropical/subtropical latitudes and are characterized by frequent cloud cover at the canopy level. They can and often do experience limited seasonality and large daily fluctuations in temperature, but not a prolonged freezing season.

The majority of taxa were coded for a single biome type, with two groups of exceptions. First, 14 living species with ranges in East and/or Southeast Asia were coded as ambiguous for warm versus cold temperate states (W/Co). Second, we assigned biome affinities to the five fossil taxa by reference to descriptions of the paleofloras from which they were obtained and paleoclimate analyses where available (e.g., Moss et al., 2005; Greenwood et al., 2010; Denk et al., 2011; Zaborac-Reed and Leopold, 2016; see Supplement 4). For four of the fossils the paleofloral assemblages imply the occupancy of forests most analogous to modern warm temperate forests. Only the paleoflora associated with the youngest fossil, an Icelandic specimen from the Mid Miocene, contained numerous cold temperate elements in addition to warm temperate species, and thus was coded as ambiguous for either warm or cold temperate (W/Co).

We simultaneously analyzed all Stage 2 *Viburnum* data—the molecular cpDNA+ITS sequences, the biogeographic data, the biome data, and the fossil data—using a Bayesian combined evidence framework (cf. Nylander et al., 2004; Pyron, 2011; Ronquist et al., 2012). Combined evidence analyses typically integrate molecular and morphological data, whereas we integrate molecular, biogeographic, and biome data. Within this framework, lineage diversification was modeled using the fossilized birth-death process (Heath et al., 2014) with an empirical stem age prior centered on 71 Ma (52.7 – 85.74 Ma; Bell & Donoghue 2005). Molecular sequences were partitioned by locus (Nylander et al., 2004) and concatenated, with each locus evolving under a GTR+Gamma substitution model (Tavare, 1986; Yang, 1994). Historical biogeography was modeled using DEC (Ree et al. 2005) with a time-stratified model (Ree & Smith 2008), with dispersal rates informed by paleogeographical dispersal graphs (Landis 2017). Biome shifts were modeled using an unconstrained rate matrix with four states. Three independent, global clocks modeled the rates of molecular substitution events, biogeographic events, and biome shift events. The molecular clock rate was further relaxed across partitions and across branches using mean-1 rate multipliers. Priors were generally chosen to be weakly informative or to place higher density on parameter values that reduce complex models into simple ones. Supplement 5 describes the model in finer detail.

When estimating the posterior, we constrained the phylogenetic relationships for three subsets of our taxa, each requiring different justifications and assumptions. Phylogenies that did not obey the clade constraints specified below were not sampled in the posterior. First, we applied a backbone constraint that required the presence of all highly supported relationships in the RAD-seq topology (i.e., clades with bootstrap support ≥ 99) in Fig. 2A.

Second, we imposed clade constraints for five fossils. Based on studies by Manchester and colleagues (e.g., Manchester, 2002; Manchester et al., 2002) we have rejected all leaf fossils previously assigned to *Viburnum*, and instead have incorporated fossil pollen grains. The pollen exine comes in three forms in *Viburnum* (Donoghue, 1985): type Ia grains are regularly reticulate, with the reticulum elevated on columellae, and with psilate (smooth) muri (Donoghue, 1985; Plates I-IV); type Ib grains have a continuous reticulum, as in Ia, but with regularly scabrate (bumpy) muri (Donoghue, 1985, Plate VI); and type Ic grains have an intectate, retipilate exine, with scabrate pili (Donoghue, 1985, Plate VII). Donoghue (1985) postulated that *Viburnum* pollen evolved from Ia to Ib to Ic, and this has been confirmed in subsequent phylogenetic analyses (e.g., Winkworth and Donoghue, 2005; Clement et al., 2011, 2014; Spriggs et al., 2015). Exine type Ib (with scabrate muri) is inferred to have evolved along the *Valvatotinus* branch, and type Ic (with scabrate pili) is inferred to have evolved independently from type Ib within the *Lentago* clade and within the *Euviburnum* clade. Two of our fossil pollen grains — one from British Columbia (Manchester et al., 2015) and one from the Florissant Fossil Beds in Colorado (Bouchal, 2013)— have the ancestral exine morphology (type Ia); these can be assigned with some confidence to *Viburnum*, and although we cannot assign them to any particular sub-clade, we forbid their assignment to sub-clades characterized by type Ib or Ic. The pollen grains from the Paris Basin in Europe (Gruas-Cavagnetto, 1978) and from the Northwest Territories in Canada (McIntyre (1991) appear to have exine type Ib, and therefore were constrained to belong to *Valvatotinus*, but were forbidden from joining the two sub-clades characterized by exine type Ic. Finally, Miocene pollen from Iceland (Denk et al., 2011) is of type Ic, and thus has two tenable topological placements. We therefore designed a specialized “either-or” clade constraint that requires the fossil placement to satisfy *either* the *Lentago*+Iceland *or* the *Euviburnum*+Iceland clade constraint.

Third, 10 of our ~163 extant *Viburnum* species are currently unsequenced. We variously constrained the placement of these species based on morphological characters. As described for fossil pollen grains, particular placements within a subclade were forbidden in several cases based on morphology. Details of these constraints are provided in Supplement 3.

The phylogenetic relationships of any unconstrained species are inferred from their molecular, biogeographic, and biome character matrices. While the five fossil taxa and the 10 unsequenced living taxa have no molecular characters, their biome and biogeography characters lend support for some phylogenetic relationships over others, and thus their phylogenetic positions are not simply drawn from the prior.

The Stage 2 analysis produces a joint posterior distribution over the constrained time-calibrated phylogenies, along with ancestral range estimates, ancestral biome estimates, and model parameters, all conditional on the combined molecular, biogeographic, biome, and fossil evidence. We apply our Stage 2 analysis in four ways: first, using a *Complete* dataset that combines all evidence currently available as described above; second, using a *Masked* dataset that sets all biogeography and biome characters for fossil taxa to be fully ambiguous (“?”); third, by recoding the four biome states into just two biome states, Non-freezing (Tropical, Cloud, Warm Temperate) and Freezing (Cold Temperate) biomes, for both the *Complete* and *Masked* datasets; and, fourth, by assessing the sensitivity of our root state estimates due to possible biome-correlated biases in fossil recovery rates (described below).

### Ancestral state estimates

Ancestral states and stochastic mappings were sampled for biogeographic ranges and biome affinities at regular intervals during MCMC sampling (Huelsenbeck et al., 2003). Each ancestral state and stochastic mapping sample corresponds to a posterior tree sample. We summarize ancestral state estimates using the maximum clade credibility (MCC) topology. Because ancestral range estimates at nodes may have differing ancestral and daughter states due to cladogenetic change, we defined a stricter node-matching rule than we used for the biome estimates, and only retained ancestral samples if the node and its two daughter nodes appear in the MCC topology.

We also present ancestral state estimates as state-frequency-through-time (SFTT) plots. An SFTT plot summarizes what proportion of lineages exist at time *t* that are found in state *i* while averaging over the posterior distribution of topologies, divergence times, and stochastic mappings. SFTT plots are generated as follows. For each posterior sample, we parse its stochastic mapping to record the number of lineages in each possible biome-area state pair (6 × 4 = 24 states) for each 1 Myr of time between 100 Ma and the Present. Within each interval, we then compute the frequency of lineages across biome states given that they are present in a particular area (biomes-per-area SFTT) and the frequency of lineages across areas given a particular biome affinity (areas-per-biome SFTT). Frequencies estimated from time bins with fewer than 510 samples are not shown, i.e. too few samples estimate all state frequencies within ±0.05 of the true multinomial frequencies with probability 0.95 (Thompson, 1987). Note that lineages without sampled descendants do not contribute to the SFTT plots. While we have achieved (to our knowledge at present) complete extant taxon sampling, our five fossil taxa almost certainly underrepresent historical *Viburnum* diversity and variation. SFTT plots were generated for both the *Complete* and *Masked* datasets to gauge the influence of fossil states on historical inferences.

### Sensitivity of biome root state estimates

All five *Viburnum* fossil pollen specimens were sampled from paleofloras that we judge to be either strictly or partially analogous to modern, warm temperate biomes, while no fossils were associated with tropical or cloud forest biomes. Because all five fossils were also sampled from North American and European localities, taphonomic or acquisition biases may distort which ancestral biomes are represented among our fossil taxa. We expect our fossil biome scores to inform our ancestral biome estimate for the most recent common ancestor of *Viburnum* (the root biome state, X_root_), but how robust would that result be to the discovery of new phylogenetic evidence? In particular, we are interested in how much “missing evidence” (e.g. from unsampled fossils) supporting alternative root states would be needed to favor a cold temperate root state. To measure this, we developed a sensitivity test that manipulates the prior probabilities for biome root state frequencies in a controlled manner. Adjusting root state priors effectively lets us specify how much missing evidence (or auxiliary data, X_aux_) we wish to introduce in favor of freezing (Co) or non-freezing (T, W, Cl) root state estimates without simulating data. We calibrate the amount of missing data by setting the cold temperate root state probability to equal 0.01, 0.02, 0.05, 0.10, 0.25, 0.40, 0.60, 0.75, 0.90, 0.95, 0.98, and 0.99, and set the root state priors for the remaining three biome states equal to (1/3) × (1 – Pr(X_root_ = Cold)). We then assess how posterior root state estimates respond to different amounts of missing evidence for both the *Complete* and *Masked* datasets.

### Biome-area transition graph

We compute the posterior mean number of times that any lineage transitioned between each ordered pair of biome-areas. Transition types that have a posterior mean less than 1 are filtered out. We then plot the posterior mean number of transitions (arrows) between biome-area pairs (nodes).

## RESULTS

### Phylogeny

Figure 2A shows the maximum clade credibility tree for the 118 *Viburnum* species for which we have RAD-seq data (branches with bootstrap support < 99 are collapsed). This tree was used as the backbone constraint tree for all Stage 2 analyses. It is almost entirely consistent with the RAD-seq tree obtained by Eaton et al. (2017) for 65 *Viburnum* species. We note, however, that the exact placement of *V. clemensiae* is unclear. Eaton et al. (2017) rooted the tree along the *V. clemensiae* branch based on previous studies using mainly cpDNA data and *Sambucus* species as outgroups (e.g., Donoghue et al., 2004; Winkworth and Donoghue, 2004; Clement et al., 2011). In the present analysis, which relied on RAD-seq data for 9 outgroup species (not shown), the position of *V. clemensiae* is equivocal and therefore collapsed. It is either sister to all other *Viburnum* species (as per Eaton et al., 2017, and previous studies), or, with equally low support, sister to a large clade containing the *Urceolata*, *Pseudotinus*, *Crenotinus*, and *Valvatotinus* clades (as in Fig. 2B). We did not recover the weakly supported (BPP=0.58) placement of *V. clemensiae* obtained by Lens et al. (2016) based on 4 cpDNA markers, ITS sequences, and outgroups (i.e., sister to a clade including all *Viburnum* except *Valvatotinus*). Other, more minor, differences in tree topology as compared to Eaton et al. (2017) and to other previous analyses (Winkworth et al., 2005; Clement et al., 2011, 2014; Spriggs et al. 2015; Lens et al., 2016) are described in Supplement 6.

Figure 2B shows the maximum clade credibility topology for the 153 species for which RAD-seq and/or chloroplast DNA sequences have been obtained (i.e., the 10 unsequenced living *Viburnum* species and the five fossils were filtered from the posterior before constructing the MCC topology). By design, this is entirely consistent with the RAD-seq backbone tree (Fig. 2A), but ~43 additional species relationships are here resolved (sometimes with poor support) based on their cpDNA and ITS sequences.

Figure 3 shows the maximum clade credibility tree for all 168 tips: 163 extant *Viburnum* species plus 5 fossils. Because we lack molecular data for 10 extant taxa and for the fossils, they behave as rogue taxa, which erodes clade support as compared to Fig. 2B (Wilkinson, 1996). That said, we see higher-than-expected support (pp=0.46 and 0.58) for the placement of two fossils (from British Columbia and Colorado) along the *Porphyrotinus* stem; this reflects their ages, presence in North America, and warm temperate biome affinities. Two other fossils (from Paris Basin and Northwest Territories) are placed along the stem of *Valvatotinus*. Our analyses favor the placement of the Iceland fossil pollen within the North American *Lentago* clade, along the *V. nudum*/*V. cassinoides* branch, rather than in the alternative position (based on exine morphology) within *Euviburnum*.

**Figure 3.**
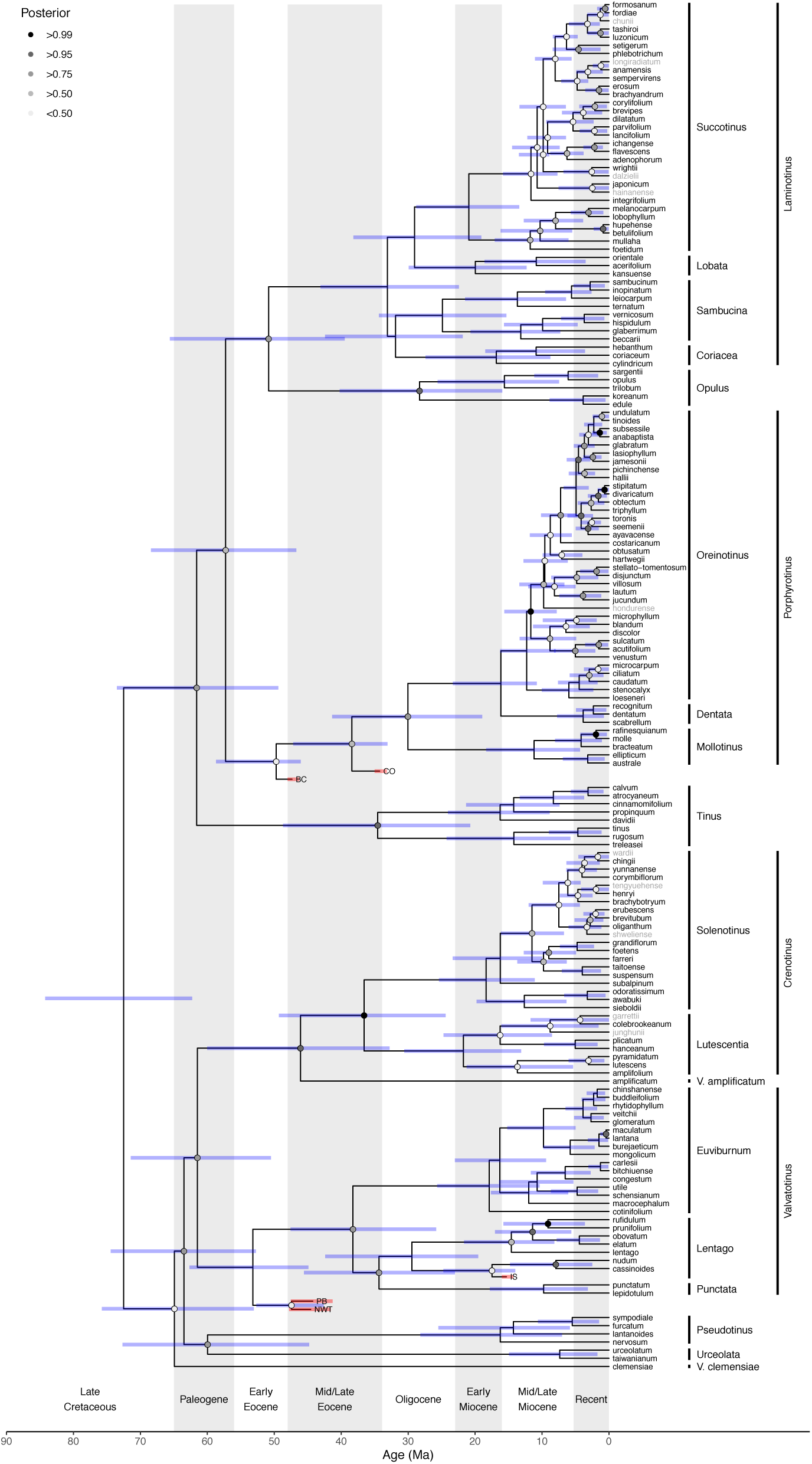
*Viburnum* divergence time estimates. Maximum clade credibility topology includes all 168 taxa from the *Complete* dataset. Node shading indicates posterior clade support. Node bars indicate 95% highest posterior density age estimates. Taxon labels for unsequenced extant taxa are colored gray.

### Dating

Figure 3 shows estimated ages (with 95% highest posterior densities, HPDs) for clades within *Viburnum*. It appears that most of the major *Viburnum* lineages had evolved by the end of the Eocene (i.e., *V. clemensiae*, *Pseudotinus*, *Urceolata*, *Punctata*, *Lentago*, *Euviburnum*, *V. amplificatum*, *Lutescentia*, *Solenotinus*, *Tinus*, *Porphyrotinus*, *Opulus*, and *Laminotinus*). The divergence of *Mollotinus* from *Dentata+Oreinotinus*, and of the *Coriacea*, *Sambucina*, *Lobata* and *Succotinus* clades, probably took place in the Oligocene. In any case, most of the diversification of *Viburnum* took place from the Miocene onward. The three significant radiations identified by Spriggs et al. (2015) – in *Succotinus*, *Oreinotinus*, and *Solenotinus* – were underway by the mid-Miocene, though most of the speciation in these clades occurred within the past 5 Myr (13/26 events in *Succotinus*, 22/34 events in *Oreinotinus*, and 8/16 events in *Solenotinus*, excluding unsequenced taxa). Overall, we estimate that 71% (120/168) of verifiable (i.e., not masked by extinction) speciation events in *Viburnum* occurred during the past 12 Myr, and 43% (72/168) within the past 5 Myr.

### Biogeography

Our reconstruction of ancestral ranges based on the full analysis is shown in Figure 4. Although the results are equivocal for the base of *Viburnum*, we note that we find the highest support for Eastern Asia alone, or for Eastern Asia plus SE Asia, and very limited support for the presence of *Viburnum* in the New World at that time. The ancestors of the several major clades that existed in the Paleocene are also inferred to have been present in Eastern Asia.

**Figure 4.**
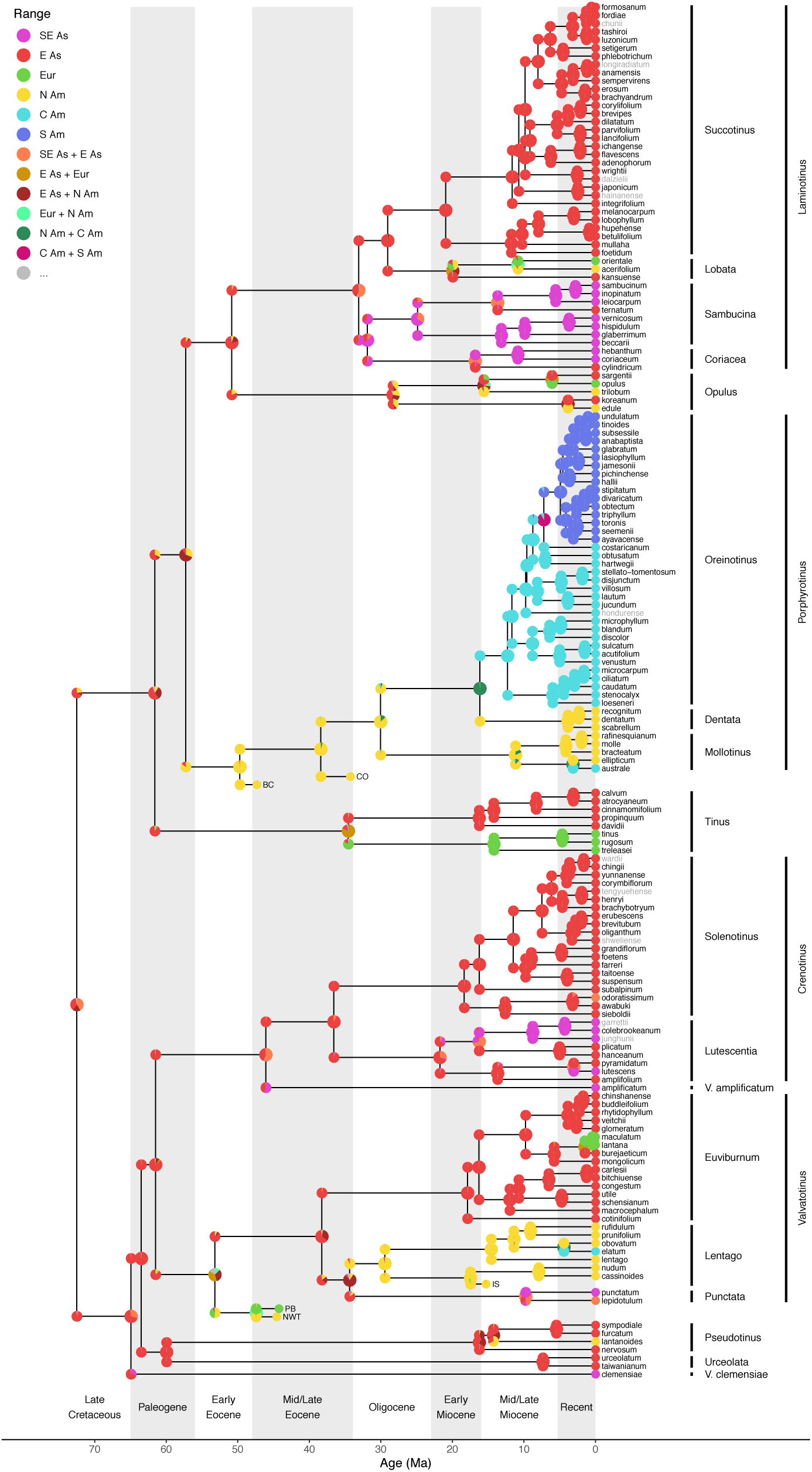
*Viburnum* ancestral range estimates taxa from the *Complete* dataset. Node colors correspond to range states (legend). Range probabilities for the ancestral range and the daughter ranges at each internal node with pie charts. Taxon labels for unsequenced extant taxa are colored gray.

By the Late Paleocene or the Early Eocene we see evidence for the movement of the *Porphyrotinus* clade into North America, most likely from eastern Asia. From North America, this lineage spread to the mountains of the Neotropics twice: (1) *V. australe* of the *Mollotinus* clade is now restricted to a few localities in northeastern Mexico; and (2) the *Oreinotinus* clade, with some 36 species, has spread as far south as Northern Argentina. The *Oreinotinus* radiation appears to have begun in eastern Mexico by the mid-Miocene and then tracked southward, with one lineage entering South America 4-10 Ma. Based primarily on the fossil pollen from the Northwest Territories, we also see the establishment of the *Valvatotinus* lineage in North America in the Eocene, although *Lentago*, the extant New World clade within *Valvatotinus*, may not have been present in North America until near the end of the Eocene. Today, *Lentago* has seven species in eastern North America and one in Mexico (*V. elatum*), extending south to Chiapas (Spriggs et al., 2019a,b).

Our results indicate that there were four other, more recent, movements into North America. The *Pseudotinus* and *Lobata* clades probably entered in the mid-Miocene, where they are represented today in eastern North America by *V. lantanoides* and *V. acerifolium*, respectively. *Viburnum lantanoides* is closely related to eastern Asian species, and we infer that its ancestor probably entered North America through the Bering Land Bridge. *Viburnum acerifolium* is sister to *V. orientale* in the Caucasus mountains of Georgia, so it is possible that it entered across the North Atlantic (Denk et al., 2011). Reconstructions for the *Opulus* clade are more complicated, but we prefer an interpretation involving two disjunctions: (1) in the core *Opulus* clade, in which the *V. trilobum* lineage may have entered North America (from Asia or Europe) perhaps as early as 15 Ma; and (2) between *V. edule* and *V. koreanum*, which today straddle the Bering strait, and may have separated within the past 4 Myr.

The rest of the diversification of *Viburnum* took place in the Old World, mainly in Eastern Asia, but with several extensions into SE Asia. Specifically, SE Asia is occupied today by the early-branching *V. clemensiae* and *V. amplificatum*, both of Borneo, the *Punctata* clade spanning from southern China to the Western Ghats of India, several species of the *Lutescentia* clade, and all members of the *Sambucina+Coriacea* clade except for *V. ternatum* (which extends into Central China).

Four clades have one or more species in Europe, and in each case these are inferred to have been derived from Asian ancestors. The *Tinus* lineage appears to have entered Europe first, perhaps at the end of the Eocene or the beginning of the Oligocene. *Viburnum tinus* is widespread around the Mediterranean (including northern Africa) and may have spawned two island endemics: *V. rugosum* in the Canary Islands and *V. treleasei* in the Azores (Moura et al., 2015). Within *Lobtata*, the ancestor of *V. orientale* may have entered Europe some 10 Ma, and the split between the European *V. opulus* and the Asian *V. sargentii* probably occurred within the past 5 Myr. Finally, in *Euviburnum* the two European species (*V. lantana* and *V. maculatum*) shared a common ancestor 1-2 Ma with their Asian relative, *V. burejaeticum*.

### Biomes

Our ancestral biome reconstructions are shown in Figure 5. With considerable certainty we infer that *Viburnum* first occupied warm temperate (lucidophyllous) forests and diversified in these forests in the Eocene. There may have been two or more shifts into more tropical (i.e., less monsoonal) forests, especially in the *V. clemensiae* and *V. amplificatum* lineages. The entry of the *Punctata* lineage and the *Coriacea+Sambucina* clade into more tropical forests probably began during the Oligocene. Support for a warm temperate origin is still favored when we mask fossil biome states (Supplement 7, Fig. S3).

**Figure 5.**
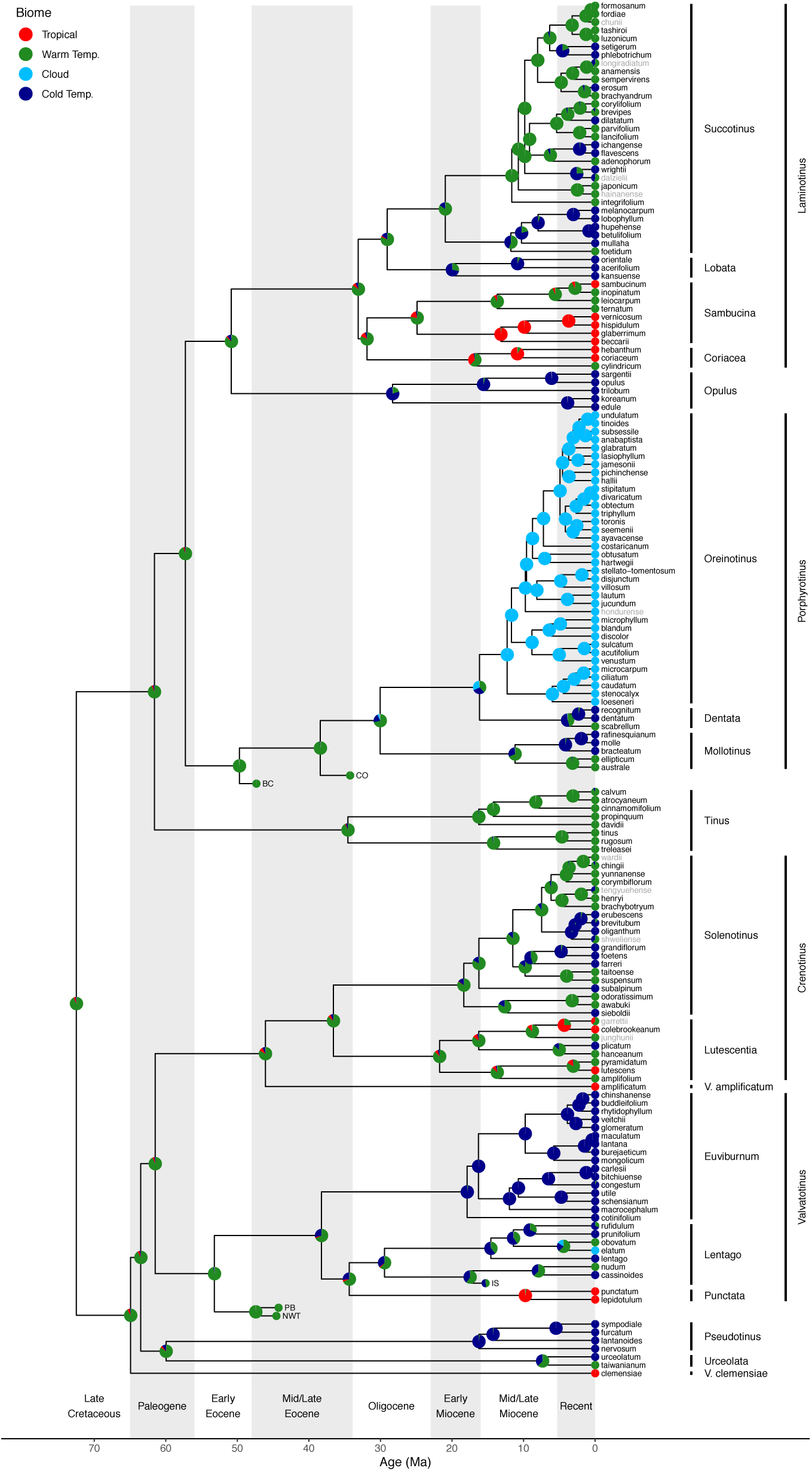
*Viburnum* ancestral biome estimates taxa from the *Complete* dataset. Node colors correspond to forest biome affinity states (legend). Probabilities are given by pie charts at nodes. Taxon labels for unsequenced extant taxa are colored gray.

There is virtually no support that *Viburnum* first diversified in cold temperate forests, regardless of whether we encode habitat preferences using four (Fig. 6) or two (Fig. S4) biomes. Instead, it appears that there were multiple transitions from warm temperate into cold temperate forests. The earliest of these shifts, perhaps in the Late Eocene, occurred in two clades (*Opulus* and *Pseudotinus*) that today occupy the coldest climates of any *Viburnum* species (Edwards et al., 2017; Park and Donoghue, in review), including boreal forests in North America (e.g., *V. edule*, *V. trilobum*, and *V. lantanoides*), Europe (V. opulus), and Asia (*V. koreanum*, *V. sargentii*). Many additional shifts into cold temperate forests appear to have taken place during the Oligocene or early Miocene. These are seen in the *Lentago*, *Euviburnum*, *Solenotinus*, *Mollotinus*, *Dentata*, *Lobata*, and *Succotinus* clades. In Asia there are multiple inferred transitions from warm to cold temperate forests within *Succotinus* (six or more) and *Solenotinus* (three or more). The only lineage that plausibly shifted from cold into warm temperate is *V. obovatum* in North America, and no lineages shifted from cold temperate into tropical biomes. It is not clear whether the Neotropical cloud forest clade (*Oreinotinus*) originated from ancestors in warm or cold temperate forests. It appears that the *Porphyrotinus* clade occupied warm temperate forests when it entered North America, and it is possible that the occupation of this biome (with limited distribution in North America today) preceded both the shift into cold temperate forests to the north and cloud forests to the south.

**Figure 6.**
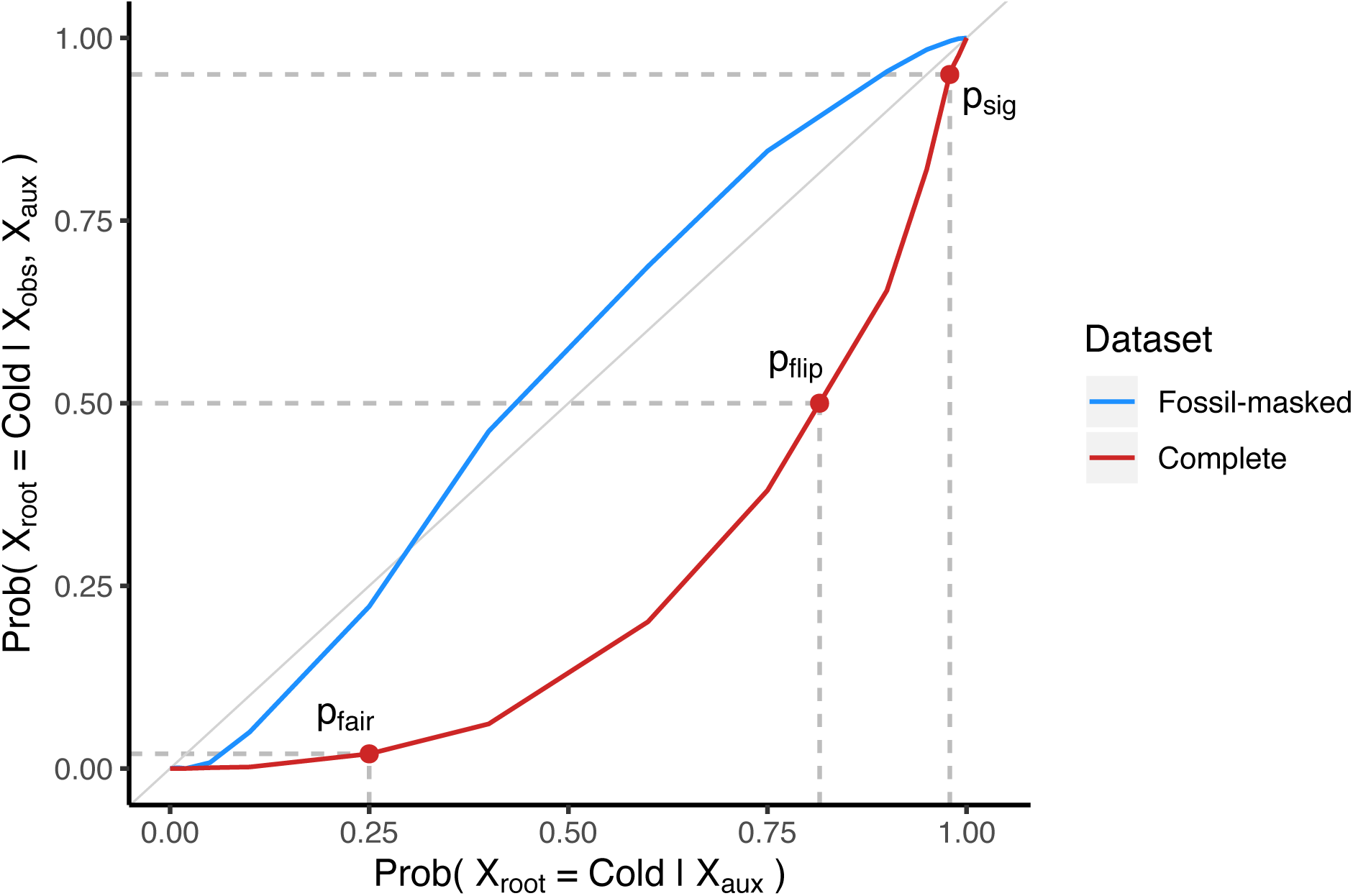
Sensitivity of *Viburnum* biome root state estimate to the addition of “missing evidence”. Root state probabilities for the cold temperate origin of *Viburnum* are estimated from the auxiliary (e.g. “missing”) data alone (x-axis) and when the observed and auxiliary data are analyzed together (y-axis). Root state probabilities both when incorporating fossil biome states (*Complete* dataset, red line) and when ignoring those states (*Masked* dataset, blue line). When the posterior mean is identical under the auxiliary data and under the observed plus auxiliary data (gray line) the observed data contain no new information about the cold root state probability. The three red markers indicate conditions where all four biome states have equal prior probability (*p*_fair_ = 0.25) and when the prior probability is sufficiently biased to estimate a cold temperate root state with posterior probability greater than 0.5 (*p*_flip_ = 0.82) or greater than 0.95 (*p*_sig_ = 0.99).

### Sensitivity Analyses

Using all four biome states, and ignoring fossil biomes with the *Masked* dataset, the posterior probability of a cold temperate biome root state is highly sensitive to the amount of missing evidence (Fig. 6). When we assign biome states to fossil taxa with the *Complete* dataset, cold temperate forests have negligible posterior support (pp < 0.05) when no missing evidence is added to the existing dataset (p_fair_ = 0.25). Cold temperate forests remain less probable than any warmer forests until the quantity of missing evidence rises to p_flip_ = 0.82, and cold temperate forests do not receive strong posterior support (pp > 0.95) until the amount of missing evidence favoring cold forests is extremely high (p_sig_ = 0.99).

### Biome and biogeography state frequencies

Reconstructed *Viburnum* lineages resided in different biomes depending on their ages and on the regions that they occupied, all of which varied over time (Fig. 7). Lineages that are 50 Ma or older generally inhabited warm temperate forests, particularly throughout the northern regions, but with no Neotropical representation. High proportions of warm temperate lineages shifted into cold forests between 35 Ma and the present, giving rise to similar proportions of warm and cold temperate lineages in Eastern Asia and Europe today, but more cold than warm temperate lineages in North America (Fig. 7, left). Early Cenozoic occupancy of cold temperate forests is far more prevalent when fossil states are *Masked* (Fig. S5, left). In SE Asia, we see low but stable proportions of tropical lineages dating back to the Eocene, with a more recent explosion of tropical lineages beginning at roughly 20 Ma (Fig. 7, left). The oldest Central American lineages (roughly 30 Ma), could have been anything but tropical, but by 15 Ma, nearly all Central American lineages are cloud forest-adapted. South American lineages are less than 10 Ma, and exclusively inhabit cloud forests.

**Figure 7.**
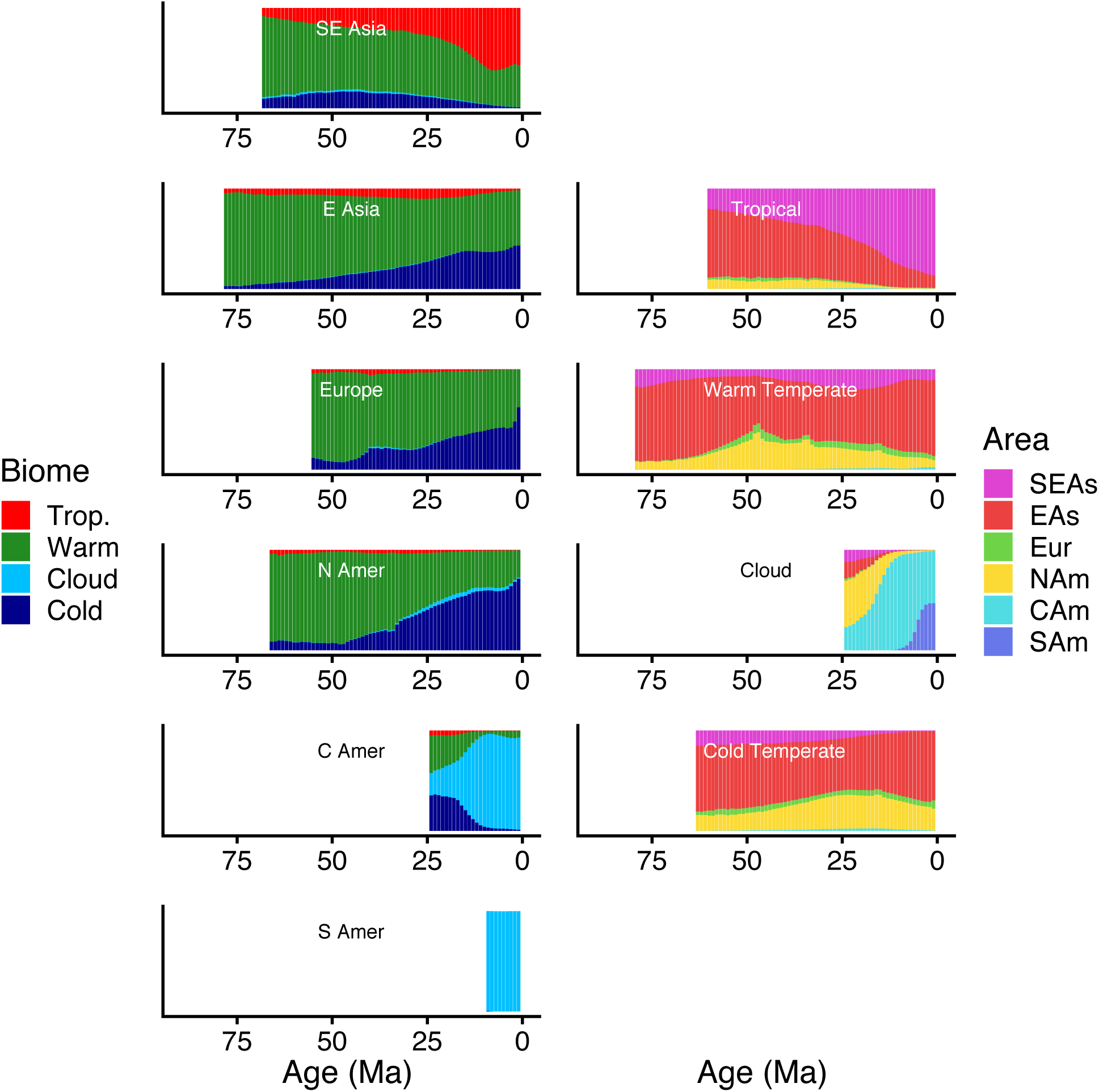
*Viburnum* biome and biogeography state frequencies through time taxa as estimated from the *Complete* dataset. Subplots in the left column report the frequency of lineages across biome states given for all lineages with sampled descendants found within a particular region. Subplots in the right column are similar, except they report regional frequencies across lineages given a particular biome state. Time bins with too few posterior samples to guarantee accurate frequency estimates were marked as empty (see main text).

Historically, most warm temperate lineages were found in East and Southeast Asia (Fig. 7, right). We find increased North American warm temperate representation that peaked at roughly 50 Ma and declined toward the present. This signal is absent when fossil biome states are *Masked* (Fig. S5, right). Although older tropical lineages were found primarily in Eastern Asia, SE Asian tropical diversity expanded steadily after 50 Ma; beginning at 20 Ma, SE Asian tropical diversity exceeds that of Eastern Asia (Fig. 7, right). The proportion of cold temperate lineages has remained fairly stable in Eastern Asia and North America over the past 50 Myr (a ratio around 2:1). Cloud forest lineages probably first appeared in North America, followed by Central America, and most recently South America.

### Biome-Area Transitions

Figure 8 shows that shifts from warm temperate into cold temperate forests have been common and have occurred most often in Eastern Asia, but also in North America. Cold-to-warm shifts are far less common in our stochastic mappings. Within the cold temperate biome, dispersals from East Asia to North America are most common. Warm temperate Asian lineages gave rise to the tropical Asian lineages. There are some transitions from Eastern Asia into SE Asian tropical forests, but few in the opposite direction. Biome shifts into Neotropical cloud forests along the branches leading to *Oreinotinus* and to *V. elatum* of the *Lentago* clade were from either cold or warm temperate forests, not from tropical forests.

**Figure 8.**
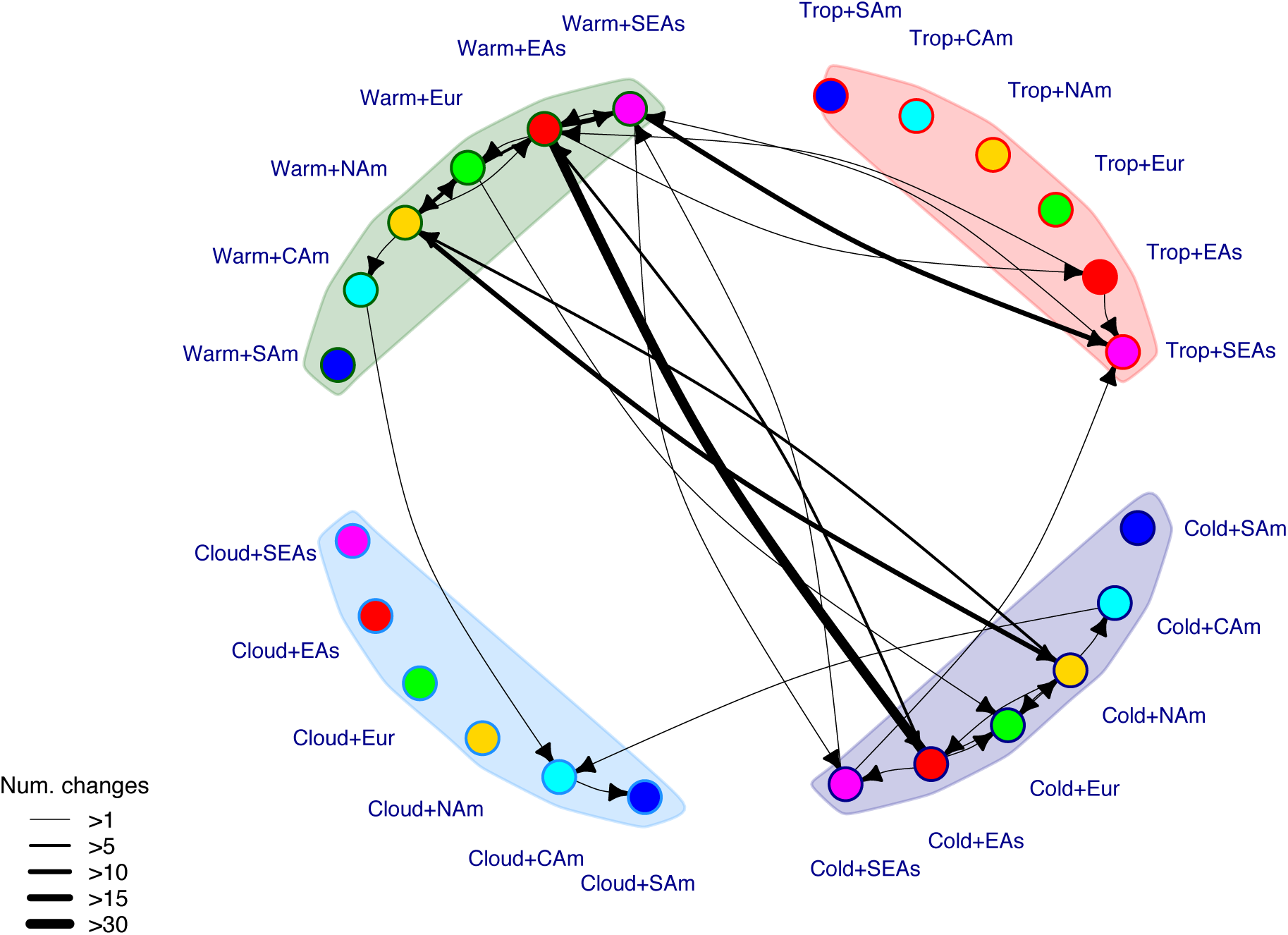
*Viburnum* biome-area transition graph estimated from the *Complete* dataset. Transparent blobs correspond to the four biomes: sub/tropical (red), warm temperate (green), cloud (cyan), cold temperate (blue) forests. Solid nodes correspond to the six regions: Southeast Asia (magenta), East Asia (red), Europe (green), North America (yellow), Central America (cyan), South America (blue). Arrows represent transitions from one region-biome state into another region-biome state. Arrows widths are classified based on the posterior mean number of transitions of the corresponding type. Transitions with posterior mean less than one are not shown.

## DISCUSSION

### Timing of Events

Several previous studies have inferred absolute ages for *Viburnum*. Using four fossils within Dipsacales, Bell and Donoghue (2005) concluded that the *Viburnum* stem lineage (i.e., the first split within crown Adoxaceae) extended into the late Cretaceous, between 70 and 85 Ma. Using fossil pollen assigned to *Valvatotinus*, Moore and Donoghue (2007, 2009) inferred a crown age for *Viburnum* in the Eocene, some 45 Ma. Using the same fossil pollen, but a greatly expanded molecular dataset, Spriggs et al. (2015) found the crown to be older, perhaps 55 Ma. Based on doubtful Eocene fossil leaves from China, Lens et al. (2016) reported a crown age for *Viburnum* centered on 56 Ma, and using disputed *Viburnum* leaves from the Paleocene pushed the crown into the late Cretaceous (Spriggs et al., 2015). Our results, based on the most complete species sample and additional fossil pollen grains, agree with a late Cretaceous estimate of over 75 Ma. Overall, increases in species sampling and additional fossils have steadily pushed back estimates of the crown age. Regardless of the exact age of the *Viburnum* crown, these analyses agree that most of the major *Viburnum* lineages were differentiated in the Eocene, and that much of *Viburnum* evolution took place from the Miocene onward.

### Biogeography

Our biogeographic results are broadly consistent with previous studies of *Viburnum*, but provide a much more complete account. Winkworth and Donoghue (2005) carried out a parsimony (DIVA; Ronquist, 1996) analysis on an undated tree for 41 species based on several chloroplast markers, nrITS, and two copies of the nuclear gene *WAXY*. They concluded that *Viburnum* originated in Asia and that there were 5-6 disjunctions between to the Old World and the New World. Most of the differences between our analyses and Winkworth and Donoghue (2005) stem from their limited species sampling. For example, because they did not include the European *V. tinus* or *V. opulus*, they identified only two disjunctions between Asia and Europe, as compared to our four.

Lens et al. (2016) analyzed *Viburnum* biogeography using a tree of 97 species based on four cpDNA markers and nrITS sequences obtained mostly from Clement et al. (2014). They reached several significantly different conclusions. Most importantly, they found several transitions from Asia to North America, followed by movements in some cases back into Asia. In contrast, our analyses identified multiple movements into North America, with no subsequent returns. These conflicts reflect differences in the sampling and phylogenetic placement of species, and the scoring of particular tips. For example, their inference that the common ancestor of the large clade that includes *Opulus* plus *Laminotinus* (i.e., *Imbricotinus*; Clement et al., 2014) was confined to North America is strongly influenced by (1) not including the Asian *V. sargentii* of the *Opulus* clade, and (2) the non-monophyly of the *Lobata* clade, and specifically the placement of the North American *V. acerifolium* as sister to rest of an otherwise Asian clade. RAD-seq data very strongly support the monophyly of *Lobata*, which effectively screens off *V. acerifolium* from influencing the ancestral area reconstruction. Consequently, we infer that the ancestors of both the *Opulus*+*Laminotinus* clade and the *Laminotinus* clade occupied Asia, and that *V. acerifolium* therefore entered North America at a later date. Similarly, our placement of *V. punctatum* (with its SE Asian sister species, *V. lepidotulum*) as sister to the *Lentago* clade (see also Eaton et al., 2017) allows us to infer that the ancestor of *Valvatotinus* occurred in Asia (rather than possibly in both North America and Asia). Finally, our finding that *V. dentatum* and *V. scabrellum* are sister to *Oreinotinus* (rather than nested within it) is consistent with previous studies and implies a much simpler biogeographic scenario (i.e., without a re-entry from the Neotropics into North America).

Leaving aside these differences, these analyses all support multiple disjunctions in *Viburnum* around the Northern Hemisphere (e.g., between Eastern Asia and Eastern North America; Eastern Asia and Europe) that broadly match those seen in many other Laurasian plant clades (Wen, 1999; Milne, 2002; Donoghue and Smith, 2004; Milne and Abbot, 2006; Wen et al., 2016). It is clear that movements through Beringia have been common on-and-off throughout the Cenozoic (Graham, 2018), and some shifts in *Viburnum* over the past 15 Myr from Asia to North America (e.g., in *Pseudotinus* and *Opulus*) are probably best explained in this way. However, in the case of *V. acerifolium*, we cannot rule out movement across the North Atlantic in this same time frame (Tiffney, 1985; Denk, 2011). Most significantly, here we are able to differentiate between four Asia-North America disjunctions that took place more recently, and two (in *Porphyrotinus* and *Valvatotinus*) that took place much earlier and may be better explained by passage across the North Atlantic (a case of pseudo-congruence; Donoghue and Moore, 2003). This difference in timing also correlates with diversification. The two clades that entered early-on are the ones that diversified in the New World, whereas the more recent arrivals are represented by just a single species. This would appear to support the thesis that the time available for diversification is a key factor in explaining diversity patterns (e.g., Wiens et al., 2011). Although one of the early New World clades, *Lentago*, contains only eight species, the other, *Oreinotinus*, has diversified rapidly in a short time period (Spriggs et al., 2015). This clade spread southward, ultimately into South America, and speciation appears to have been elevated owing to isolation in disjunct Neotropical mountain regions. In contrast, the rapid radiation of the *Succotinus* clade in Asia (Spriggs et al., 2015) appears to have been fostered by disjunctions involving China, Korea, Japan, and Taiwan, coupled with multiple shifts from warmer to colder forests.

### Ancestral biomes and biome shifts

Previous analyses have reached opposite conclusions regarding the biome in which *Viburnum* initially diversified. Spriggs et al. (2015) supported the view that *Viburnum* originated in tropical forests, whereas Lens et at. (2016) concluded that it evolved in cold temperate forests. Our analyses strongly support a third interpretation, namely that *Viburnum* originated in warm temperate (lucidophyllous) forests, and subsequently adapted to both colder and to more tropical climates (Fig. 5). When we collapsed the comparison to warmer forests that experience little freezing versus cold forests with prolonged freezing, we very confidently favored an origin in the warmer forests, and decisively ruled out cold temperate forests as the original biome (Fig. S4). However, with respect to distinguishing warm temperate versus tropical forests we caution that our analyses do not address the possibility that there have been differential rates of extinction in the different biomes. Specifically, they do not test the hypothesis advanced by Spriggs et al. (2015) that rates of extinction in *Viburnum* have been higher in tropical forests since the Oligocene. If this was indeed the case, our modern sample would bias against obtaining a tropical ancestry.

Convincing estimation of ancestral biomes is tricky, especially in light of extinction, but it benefits from the comprehensive sampling achieved here and insights provided by all relevant data sources. Specifically, we highlight that the inclusion of fossil biome assignments has a major impact. Warm temperate forests are favored as ancestral whether the fossils are included (Fig. 5) or not (Fig. S3), but their inclusion does dramatically increase our confidence in this result. Indeed, as our sensitivity analyses show (see above), a substantial increase of “missing evidence” would be needed to erase the influence of known fossil biome states, and even more to achieve significant support for cold forest ancestry.

Regarding subsequent biome shifts, it appears that there were multiple shifts into more tropical climates from warm temperate regions, some possibly quite early in *Viburnum* evolution (*V. clemensiae*, *V. amplificatum*, the Punctata clade, the *Sambucina*+*Coriacea* clade, and several *Lutescentia* species). Of these, we note the tropical radiation of the newly recognized *Sambucina*+*Coriacea* clade, within which there have been a number of recent speciation events involving disjunctions between the Philippines (*V. glaberrimum*), Borneo (*V. vernicosum* and *V. hispidulum*), and Peninsular Malaysia (*V. beccarii*).

Likewise, there have been multiple shifts from warm temperate into cold temperate forests, scattered throughout the tree: e.g., the *Opulus* and *Pseudotinus* clades, *V. plicatum* within *Lutescentia*, *V. grandiflorum* and its relatives within *Solenotinus*, and multiple instances within *Succotinus* (Fig. 5). The largest single radiation in cold temperate forests is *Euviburnum* with 16 species, mostly in central China, but extending along the Himalayas (*V. cotinifolium*) and into Europe (*V. lantana* and *V. maculatum*). Importantly, we inferred no clear-cut reversals from cold forests back into warm or tropical forests.

### The Interaction of Geographic Movements and Biome Shifts

The combined analysis of biogeography, biomes, and fossils provides us with several new insights. The premier case concerns when, and in which biome, the large *Porphyrotinus* clade entered the New World. Our combined analyses unequivocally support the view that *Porphyrotinus* arrived in the New World in the Early Eocene. This reflects the presence of *Viburnum* fossil pollen in the New World in the mid-Eocene, and the placement of the grains from British Columbia (BC) and Colorado (CO) with moderate support along the stem of *Porphyrotinus* (BC+CO+*Porphyrotinus*, pp=0.47; CO+*Porphyrotinus*, pp=0.59; BC+*Porphyrotinus*, p=0.16). It is likely their shared geography that unites these fossils with *Porphyrotinus*, as support for these relationships plummet when fossil states are masked (BC+CO+*Porphyrotinus*, pp=0.02; CO+*Porphyrotinus*, pp=0.14; BC+*Porphyrotinus*, pp=0.10). Alternative placements of these pollen grains with East Asian clades would require additional (less probable) dispersal events.

Most importantly, our results suggest that the *Porphyrotinus* lineage originally occupied warm temperate forests. This is significant for two reasons. First, it implies that there were later shifts into cold temperate forests within North America. That is, these plants adapted *in situ* to the spread of cold climates, as opposed to having arrived from Asia already adapted to them. Second, it implies that warm temperate forests were widespread in the Eocene in North America, whereas today these forests are almost absent from the region except in the SE United States. Less integrated analyses would suggest instead that *Porphyrotinus* entered the New World already adapted to cold forests. Likewise, our results allow cloud forest plants to have evolve directly from warm temperate plants, whereas less complete analyses would favor their origin from cold temperate plants.

In the case of the lineages that arrived in the New World more recently, our results indicate that these had already adapted to cold climates in the Old World. For example, the eastern North American *V. lantanoides* is nested within an otherwise Asian clade (*Pseudotinus*), and its immediate ancestor was presumably already adapted to cold when it spread (Park and Donoghue, 2019). The same logic applies to *V. acerifolium* in *Lobata*, and to intercontinental movements in the two subclades within *Opulus*. Overall, as expected given global cooling since the Eocene, the *Viburnum* occupancy of cold temperate forests has increased through time in North America, as it also has in Asia (owing to multiple evolutionary shifts from warm to cold forests) and in Europe (owing to migration) (Figs. 7, 8). Shifts into tropical forests have occurred in Asia, but never elsewhere. In particular, we note that there have been no transitions into tropical forests from cloud forests in the Neotropics, despite ample opportunity based on the direct adjacency of these biomes throughout that region (Donoghue and Edwards, 2014).

### The Evolution of Biome-Related Traits

Lens et al. (2016) compared *Viburnum* with its relative *Sambucus* in order to test the theory (see Carlquist, 1975, 2012) that simple perforations in vessel elements in the wood evolved from scalariform perforations in connection with a shift into dry and/or warm areas where there would be selection for greater hydraulic efficiency. They documented scalariform perforations in 17 species of *Viburnum* and simple perforations in 17 species of *Sambucus*, and they assessed the contemporary and inferred ancestral climate spaces occupied by these two clades. They found that the modern climates occupied by a selection of *Viburnum* and *Sambucus* species are largely overlapping, but they argued that *Viburnum* evolved initially in colder climates and *Sambucus* in warmer ones. Specifically, they argued that *Viburnum* likely evolved in cold temperate forests, and that freezing temperatures in these forests explain why scalariform perforations were retained (i.e., to prevent freezing-induced air embolisms).

In stark contrast, we find that *Viburnum* radiated initially in warm temperate forests that experienced little freezing, and moved multiple times into colder forests. As detailed in Supplement 6, this major difference in our two results stems from their limited species sampling and inaccurate climate scores for several key species. Our findings imply that exposure to cold was *not* the factor that favored the retention of scalariform perforations. It is plausible, however, that they retained ancestral vessel elements simply because they continued to occupy mesic forests (albeit without prolonged freezing) where they did not experience a significant increase in evaporative demand (Carlquist, 1975, 2012). This does not negate their argument for the evolution of simple perforations in *Sambucus* (we have not reanalyzed whether this lineage shifted into warmer/drier areas requiring greater hydraulic efficiency), but it does alter their conclusions regarding *Viburnum* evolution and the presumed benefits of scalariform perforations.

Regarding the evolution of other biome-relevant traits, we note that our results broadly support the conclusions of other studies regarding leaf traits. Specifically, our analyses are consistent with the view that the first viburnums had more-or-less entire (smooth) or irregularly toothed leaf margins, and that shifts into colder climates in multiple lineages were accompanied by the evolution of regularly toothed and/or lobed leaves (Schmerler et al., 2012; Spriggs et al., 2018; Edwards et al., 2016). Likewise, our findings support multiple shifts from the evergreen habit to the deciduous habit associated with adaptation to climates with prolonged winter freezing (Edwards et al., 2017). We note that there have also been reversals in both characters; for example, evergreen leaves with entire margins evolved in the cloud forests of the Neotropics. Overall, the concordance of biome shifts with biome-related traits reinforces the hypothesis that *Viburnum* originated in warm forests.

## CONCLUSIONS

We draw several general lessons from our analyses. First, as advocates for comprehensive sampling (Donoghue and Edwards, 2019), we have done our best to include all of the *Viburnum* species that we currently recognize. Yet, it is important to appreciate that we still come up short. First, basic taxonomic issues remain, and we anticipate the discovery of additional *Viburnum* species as we apply molecular approaches at the population level. A case in point is our resurrection of *V. nitidum*, which was hidden on the coastal plain of the Southeastern US (Spriggs et al., 2019). Second, we are painfully aware of how little we know about extinct viburnums. The structure of the tree itself strongly suggests that we are missing such species. Tropical depauperons such as *V. clemensiae* and *V. amplificatum* imply past extinctions (Donoghue and Sanderson, 2015), and, as Spriggs et al. (2015) argued, rates of extinction in *Viburnum* may have been especially high in the tropics. What appear to be abrupt major biome transitions (e.g., directly from warm forests into boreal forests in *Pseudotinus* and *Opulus*) also suggest that we are missing ecological intermediates. The failure to fully represent these lineages surely biases our studies.

Utopians might argue that we will eventually obtain the perfect dataset for all species of interest. But before then, biologists must integrate diverse datasets, like ours, that vary in terms of species sampling, character completeness, data quality, biological properties, and ease of modeling. The integration of these data presents challenges that we suspect others face, too. We hope that our approach will be of some general interest. During our preliminary analyses, for instance, we found that our newer RAD-seq dataset was reliable for estimating tree topology (Eaton et al., 2017), but the large numbers of short loci were not easily modeled using standard divergence time estimation methods. The longer loci from our species-rich cpDNA dataset, on the other hand, could be used to estimate node ages, even if cpDNA alone is not ideal for estimating species relationships at all timescales. These data had complementary strengths for estimating the evolution of *Viburnum*, so no hard-won data were discarded. Moreover, reusing our older (but here expanded) cpDNA dataset allowed us to draw stronger conclusions regarding the effect of the data on the inference (Supplement 6). Like the molecular datasets, incorporating the fossil, biome, and biogeography datasets each presented its own challenges, which we discuss below.

In *Viburnum*, as in most other clades, critically identified fossils remain rare. Fortunately, in this case there are a few fossils, and we have tried to make the most of the limited evidence that these provide. For example, we developed an “either-or” clade constraint to ensure that our Icelandic fossil formed a clade with either of two non-sister extant clades. Without this constraint, we would have needed to force a relationship with *Euviburnum* or with *Lentago*, or to relax the constraint to allow the unfounded placement of the fossils among the lineages that separate *Euviburnum* from *Lentago*, or to neglect the fossil altogether. In connection with this development, we added functionality to RevBayes to apply multiple backbone constraints to overlapping taxon sets, which was necessary to infer the relationships of the morphology-only species.

Including fossil taxa with their geographic and biome states has two effects: it helps affiliate a fossil taxon with particular extant clades, and it tethers the ancestral condition of these clades through that affiliation. As discussed above, we saw this especially in the case of the *Porphyrotinus* clade. If, instead, we had adopted a sequential inference strategy – first estimating the tree distribution and secondarily estimating ancestral states across that distribution – we would likely have placed the British Columbia and Colorado fossils apart from *Porphyrotinus* in roughly 75% of sampled phylogenies, which would proportionally reduce support for *Porphyrotinus* entering North America during the Eocene. These rogue fossils would in turn diffusely inflate support for North American and warm temperate ancestry throughout the tree. This is avoided in our joint inference framework, illustrating that biogeography has another promising role to play in phylogenetic inference (cf. Landis, 2017, Landis et al., 2018).

On the other hand, we recognize that there are potential biases in the sampling of fossils. Our fossil pollen was obtained from predominantly warm temperate plant assemblages in North American and European localities, and none were sampled from East Asia, where *Viburnum* probably originated. By treating the prior root state frequencies as a surrogate for missing root state evidence, we developed a simple Bayesian sensitivity test to measure what quantity of newly discovered evidence would be needed to recover a cold temperate ancestral state estimate (Fig. 6). This revealed that we would require new evidence equivalent to a 3-fold increase in prior weight for freezing forests to attain majority support (pp > 0.5) for a freezing root state. One way to interpret this result is that it would require the discovery of twelve new cold temperate fossils from the Paleocene or Eocene (i.e., three times the four warm temperate fossil pollen samples that we have from those periods), while excluding any new discoveries of fossils from non-freezing biomes. Overall, our sensitivity test echoes earlier findings that taxon sampling really matters (Heath et al., 2008, Wright et al. 2015, Matzke et al. 2012, Betancur et al. 2015). Future sensitivity studies might complement the existing extant and fossil taxa with simulated fossil taxa under a panel of geography- and biome-dependent biases.

We are also keenly aware of the need to propagate uncertainty throughout the joint Bayesian inference of evolutionary history. Among other things, this creates new challenges for visualization. Ancestral state trees are typically plotted against a single (consensus) phylogeny (Figs. 3 and 4). We complemented this simplified view of range and biome evolution with plots of state frequencies through time that marginalized over various sources of uncertainty, including topology among fossil and extant taxa, divergence times, stochastic mappings, and evolutionary parameters. Rather than generating M x N = 24 plots for all pairs of M = 4 biomes and N = 6 regions, we instead developed M + N = 10 conditional plots to highlight how the occupancy of biomes and regions co-evolved in *Viburnum*. By averaging over uncertainty in this way, we gain a clearer summary of evolutionary trends independent of any specific phylogenetic hypothesis. For example, we are able to conclude that the proportion of cold temperate viburnums exceeded 25% in East Asia, Europe, and North America starting in the Miocene (Fig. 7).

The combined evidence approach developed here has allowed us to integrate molecular data, fossils, biogeographic ranges, and biome affinities, and this has provided a new and greatly improved understanding of the geographic and ecological radiation of a plant clade through the Cenozoic. But, there are clearly other relevant lines of evidence, and we look forward to their incorporation as well. For example, in addition to the leaf and wood traits mentioned above, we are optimistic about more fully and directly incorporating physiological and phenological data for *Viburnum* that bear on biome occupancy (e.g., Chatelet et al., 2013; Edwards et al., 2014; Scaffoni et al., 2016). Recent work to estimate ancestral biome affinities within an evolving spatial distribution of biomes through time also speak to the warm or cold origin of *Viburnum* (Landis et al., in review). In the end, it is the alignment of these accumulating lines of evidence that lends confidence that we are on the right track.

## SUPPLEMENTARY MATERIAL

Supplementary Information is organized into the following sections within the accompanying document: (1) RAD-seq data, (2) Chloroplast DNA data, (3) Placement of unsequenced species based on morphological traits, (4) Fossil pollen morphology and biome assignments, (5) Bayesian phylogenetic analysis, (6) Extended discussion of phylogenetic and ancestral biome estimates, and (7) Supplementary Figures (S1 through S8). Raw RAD-seq data have been submitted to NCBI SRA under project number PRJNAXXX. All remaining data are available from the Dryad Digital Repository: http://dx.doi.org/10.5061/dryad.####.

## ACKNOWLEDGEMENTS

Our studies have relied heavily on specimens housed in many herbaria, large and small. We are extremely grateful for the careful stewardship (and, increasingly, the digitization) of these priceless collections, and to the many collectors of these specimens over the centuries. Our work has also built on our field studies in over a dozen countries in the past two decades (Vietnam, Malaysia, Indonesia, the Philippines, China, Taiwan, Japan, India, Georgia, Colombia, Ecuador, Peru, Bolivia, and Mexico), and we are forever indebted to the many colleagues who have made these trips possible and so much fun. Finally, we wish to express our gratitude to the staffs of the Arnold Arboretum of Harvard University and the Harvard University Herbaria, where we have spent countless hours studying our favorite plants.

