## Supplementary Information for "Joint phylogenetic estimation of geographic movements and biome shifts during the global diversification of *Viburnum*"

**Supplement 1. RAD-seq data**

Table S1 identifies the 127 species and vouchers used to generate the RAD-seq dataset. Of the 127 species, 1 species is from *Adoxa*, 1 is from *Sinadoxa*, 7 are from *Sambucus*, and the remaining 118 species belong to *Viburnum*. Raw RAD-seq data have been submitted to NCBI SRA under project number PRJNAXXX.

Collector names are coded as BP = Brian Park, DARE = Deren Eaton, DB = David Boufford, DE = Donald Egolf, DM = David Michner, DS = Douglas Stevens, EE = Erika Edwards, ES = Elizabeth Spriggs, F = Fuentes, KFC = Kao-Fang Chung, MD = Michael Donoghue, PS = Patrick Sweeney, RW = Richard Winkworth, SFD = Sabah Forestry Department, T = Takahashi, TE = Torsten Eriksson, TY = Tetsukasu Yahara, WA = William Alverson, and WC = Wendy Clement. Herbarium names are coded as A = Arnold Arboretum, Harvard University Herbaria, BRU = Brown University Herbarium, GH = Grey Herbarium, Harvard University, S = Swedish Museum of Natural History, USNA = US National Arboretum, WIS = Wisconsin State Herbarium, and YU = Yale University Herbarium. Specimens without precise voucher information are coded under "Herbarium Code" as VL = Voucher lacking, VM = Voucher missing, and VU = Voucher unknown. Voucher lacking (VL) indicates that no voucher was ever collected for the specimen. Voucher missing (VM) indicates that we believe a voucher was collected, but we have not obtained it yet. Voucher unknown (VU) means that we are unaware if a voucher actually exists for a given specimen in an herbarium.

| Genus | Species | Collector Code | Collection Number | Herbarium Code |
|---|---|---|---|---|
| *Adoxa* | *moschatellina* | MD | 1 | WIS |
| *Sinadoxa* | *corydalifolia* | DB | 26555 | A |
| *Sambucus* | *canadensis* | ES | 242 | YU |
| | *callicarpa* | Unknown | cult. Rancho Santa Ana #14770 | VU |
| | *javanica* | T | 1783 | A |
| | *nigra* | TE | s.n. | S |
| | *racemosa* | TE | s.n. | S |
| | *sieboldiana* | Unknown | cult. AA 1887-77-G | A |
| | *williamsii* | DB | 27670 | A |
| *Viburnum* | *acerifolium* | ES | 88 | YU |
| | *acutifolium* | MD | "Mirador", Highway 175, Oaxaca, Mex. November 6, 2005 | VL |
| | *amplificatum* | SFD | 156003 | YU |
| | *amplifolium* | PS | 2252 | YU |
| | *anamensis* | PS | 2094 | YU |
| | *awabucki* | DE | 2344E | USNA |
| | *beccarii* | PS | 2084 | YU |
| | *betulifolium* | KFC | 1950 | YU |
| | *bitchiuense* | Unknown | cult. AA 2047-77B | A |
| | *brachyandrum* | MD | MJD JP-10 | YU |
| | *brachybotryum* | PS | 2222 | YU |
| | *buddleifolium* | PS | 2607 | YU |
| | *carlesii* | BP | BP-001 | YU |
| | *caudatum* | MD | C2 | YU |
| | *chinshanense* | PS | 2269 | YU |
| | *ciliatum* | PS | 3220 | YU |
| | *cinnamomifolium* | PS | 2105 | YU |
| | *clemensiae* | PS | 2135 | YU |
| | *congestum* | PS | 2235 | YU |
| | *coriaceum* | PS | 2089 | YU |
| | *coryfolium* | PS | 2249 | YU |
| | *costaricanum* | MD | 85 | YU |
| | *cotinifolium* | WC | 267 | YU |
| | *cylindricum* | WC | 268 | YU |

| | | | | |
|---|---|---|---|---|
| | *davidii* | WC | 269 | YU |
| | *dentatum* | ES | 650 | YU |
| | *dilatatum* | PS | 2212 | YU |
| | *disjunctum* | MD | MJD-4, 1989 (MJD66-2) | YU |
| | *divaricatum* | PS | 1773 | YU |
| | *edule* | WA | Northern Wisconsin (MJD56) | VL |
| | *elatum* | PS | 3084 | YU |
| | *ellipticum* | MD | s.n., vic. Corvallis, Oregon (MJD49)<br>July 7, 2012 | YU |
| | *erosum* | PS | 2220 | YU |
| | *erubescens* | RW | 36 | YU |
| | *farreri* | RW | 21 | YU |
| | *foetens* | WC | 270 | YU |
| | *foetidum* | PS | 2250 | YU |
| | *fordiae* | PS | 2201 | YU |
| | *formosanum* | PS | 2202 | YU |
| | *furcatum* | ES | 735 | YU |
| | *glaberrimum* | PS | 2323 | YU |
| | *glomeratum* | PS | 2559 | YU |
| | *grandiflorum* | WC | 271 | YU |
| | *hallii* | PS | 1830 | YU |
| | *hanceanum* | PS | 2195 | YU |
| | *hartwegii* | PS | 3190 | YU |
| | *henryi* | WC | 272 | YU |
| | *hispidulum* | PS | 2136 | YU |
| | *ichangense* | DM | 1160 (cult. AA 51-60-A) | A |
| | *inoptinatum* | PS | 2091 | YU |
| | *integrifolium* | KFC | 1946 | YU |
| | *jamesonii* | PS | 1636 | YU |
| | *japonicum* | WC | 272 | YU |
| | *jucundum* | PS | 3188 | YU |
| | *kansuense* | DB | 27416 | A |
| | *lantana* | DM | 680 (cult. AA 1089-60A) | A |
| | *lantanoides* | ES | 70 | YU |
| | *lautum* | PS | 3189 | YU |
| | *leiocarpum* | PS | 2265 | YU |
| | *lentago* | ES | 401 | YU |

| | | | | |
|---|---|---|---|---|
| | *lobophyllum* | MD | 25 | YU |
| | *loesnerii* | DS | 2547 | YU |
| | *lutescens* | PS | 2077 | YU |
| | *luzonicum* | KFC | 1947 | YU |
| | *macrocephalum* | WC | 284 | YU |
| | *microcarpum* | PS | 3202 | YU |
| | *microphyllum* | DARE | 3 | YU |
| | *molle* | MD | 5 | YU |
| | *muhalla* | WC | 274 | YU |
| | *nervosum* | PS | 2288 | YU |
| | *nudum* | ES | 309 | YU |
| | *nudum* | ES | 621 | YU |
| | *obovatum* | ES | 595 | YU |
| | *obtusatum* | PS | 3100 | YU |
| | *odoratissimum* | EE | 2013-35 | BRU |
| | *opulus* | WC | 250 | YU |
| | *orientale* | MD | s.n., Georgia | YU |
| | *parvifolium* | KFC | 1953 | YU |
| | *pastasanum* | PS | 1799 | YU |
| | *phlebotrichum* | ES | 774 | YU |
| | *pichinchense* | PS | 1665 | YU |
| | *pichinchense* | PS | 1669 | YU |
| | *plicatum* | ES | 723 | YU |
| | *propinquum* | PS | 2596 | YU |
| | *prunifolium* | ES | 478 | YU |
| | *punctatum* | PS | 2097 | YU |
| | *pyramidatum* | PS | 2272 | YU |
| | *recognitum* | ES | 378 | YU |
| | *reticulatum* | PS | 1702 | YU |
| | *reticulatum* | PS | 1719 | YU |
| | *rhytidiphyllum* | Unknown | cult. AA 1386-82B | A |
| | *rufidulum* | ES | 221 | YU |
| | *sambucina* | PS | 2100 | YU |
| | *sargentii* | ES | 724 | YU |
| | *scabrellum* | ES | 193 | YU |
| | *schensianum* | PS | 2565 | YU |
| | *seemenii* | F | 4724 | GH |
| | *sempervirens* | PS | 2191 | YU |
| | *setigerum* | Unknown | cult. AA 305-2002A | A |

| | | | | |
|---|---|---|---|---|
| | *sieboldii* | ES | 721 | YU |
| | *stellato-tomentosum* | MD | 640 | YU |
| | *stenocalyx* | PS | 3208 | YU |
| | *subalpinum* | PS | 2292 | YU |
| | *sulcatum* | MD | "Mirador", Highway 175, Oaxaca, Mex. November 6, 2005 | VL |
| | *suspensum* | MD | 36 | YU |
| | *sympodiale* | KFC | 1932 | YU |
| | *taitoense* | PS | 2229 | YU |
| | *taiwanianum* | KFC | 1952 | YU |
| | *tashiori* | TY | Amami Island, Japan November 18, 2011 | VM |
| | *ternatum* | PS | 2264 | YU |
| | *tinus* | WC | 277 | YU |
| | *trilobum* | RW | 14 | YU |
| | *triphyllum* | EE | 2014-04 | YU |
| | *urceolatum* | MD | MJD JP-8 | YU |
| | *utile* | PS | 2593 | YU |
| | *veitchii* | Unknown | cult. AA 457-94B | A |
| | *vernicosum* | PS | 2123 | YU |
| | *wrightii* | MD | MJD JP-7 | YU |

Table S1. RAD-seq vouchers.

**Supplement 2. Chloroplast DNA data**

This section reports voucher information and GenBank accession data for 157 *Viburnum* species included in this phylogenetic study. Much of the data were published in Clement and Donoghue (2011), Chatelet et al. (2013), and Clement et al. 2014. Species new to the study of *Viburnum* phylogeny are indicated in bold and underlined text; species that have previously been included in the *Viburnum* phylogeny but have been replaced with accessions from field collections (P.W. Sweeney, M.J. Donoghue, W.L. Clement, and E.J. Edwards) are indicated in bold. Species are organized by clade following the phylogenetic nomenclature published by Clement et al. (2014). Herbarium abbreviations are as follows: A = Arnold Arboretum, Harvard University Herbaria, GH = Grey Herbarium, Harvard University, K = Kew Royal Botanic Gardens, MO = Missouri Botanical Garden, NY = New York Botanical Garden, WTU = University of Washington Herbarium, and YU = Yale University Herbarium. A hyphen (-) in the list of GenBank accessions indicates missing data.

Clade. Taxon; voucher specimen (herbarium). ITS; *mat*K; *ndh*F; *pet*B-*pet*D; *rbc*L; *rpl*32-*trn*L$^{(UAG)}$; *trn*C-*ycf*6; *trn*H-*psb*A; *trn*K; *trn*S-*trn*G.

***Viburnum***. *Viburnum clemensiae* J. Kern: J. Beaman 11781 (K). AY265117; HQ591569; HQ591648; HQ591999; HQ591714; HQ591878; HQ592122; AY627387; AY265163; EF490267.

***Regulaviburnum***.

***Valvatotinus***.

***Lentago.*** *Viburnum cassinoides* L.: E.L. Spriggs ELS79 (YU). pending; pending; pending; pending; pending; pending; pending; pending; -; pending. *Viburnum elatum* Benth.: P.W. Sweeney et al. PWS3063 (YU). pending; pending; pending; pending; pending; pending; -; pending; -; pending. *Viburnum lentago* L.: M.J. Donoghue & R.C. Winkworth 21 (YU). AY265136; HQ591598; HQ591670; HQ592022; HQ591739; HQ591905; HQ592148; AY627406; AY265182; EF490280. *Viburnum nudum* L.: E.L. Spriggs ELS29 (YU). pending; pending; pending; pending; pending; pending; pending; pending; pending; pending. *Viburnum obovatum* Walter: E.L. Spriggs ELS264 (YU). pending; pending; pending; pending; pending; pending; pending; pending; pending; pending. *Viburnum prunifolium* L.: M.J. Donoghue & R.C. Winkworth 13 (YU). AY265144; HQ591615; HQ591683; HQ592033; HQ591756; HQ591922; HQ592163; AY627413; AY265190; EF490286. *Viburnum rufidulum* Raf.: M.J. Donoghue & R.C. Winkworth 14 (YU). AY265147; HQ591620; HQ591687; HQ592038; HQ591761; HQ591927; HQ592167; AY627415 ; AY265193; EF490287.

***Paleovaltinus***

***Punctata.*** *Viburnum lepidotulum* Merr. & Chun: P.W. Sweeney et al. PWS2097 (YU). KJ795805; KF019748; KF019768; KF019825; KF019790; KF019866; KF019889; -; KF019934; KF019913. *Viburnum punctatum* Buch.-Ham. ex D.Don: P.W. Sweeney et al. PWS2274 (YU). -; pending; pending; pending; pending; pending; pending; pending; -; pending.

***Euviburnum.*** *Viburnum bitchiuense* Makino: David Chatelet 1097-77A (A-living collection). JX049448; JX049451; JX049459; JX049509; JX049471; JX049477; JX049481; JX049467; JX049491; JX049495. *Viburnum buddleifolium* C.H. Wright: P.W. Sweeney et al. PWS2607 (YU). pending; pending; pending; pending; pending; pending; pending; pending; pending; pending. *Viburnum burejaeticum* Regel & Herder: K. Schmandt 375-95A, 00223095 (A). -; JQ805231; JX049463; JX049513; JX049463, JX049473; JQ805472; JX049486; JQ805297; JQ805552; JX049500. *Viburnum carlesii* Hemsl. ex Forb. & Hemsl.: M.J. Donoghue & R.C. Winkworth 24 (YU). AY265115; HQ591566; HQ591645; HQ591996; HQ591710; HQ591873; HQ592117; AY627385; AY265161; HQ591823. ***Viburnum chinshanense*** **Graebn.: P.W. Sweeney et al. PWS2269 (YU). pending; pending; pending; pending; pending; pending; pending; pending; pending; pending.** *Viburnum congestum* Rehder: P.W. Sweeney et al. PWS2235 (YU). pending; pending; pending; pending; pending; pending; pending; pending; pending; pending. *Viburnum cotinifolium* D. Don: M.J. Donoghue WC267 (YU). KF019809; KF019744; KF019767; KF019823; KF019787; KF019864; -; KF019843; KF019932; KF019908. *Viburnum glomeratum* Maxim.: P.W. Sweeney et al. PWS2559 (YU). pending; pending; pending; pending; pending; pending; pending; pending; pending; pending. *Viburnum lantana* L.: M.J. Donoghue & R.C. Winkworth 26 (YU). AY265134; HQ591595; HQ591667; HQ592019; HQ591736; HQ591902; HQ592145; AY627404; AY265180; EF490278. *Viburnum macrocephalum* Fortune: M.J. Donoghue 101 (YU).

EF462984; HQ591604; HQ591673; HQ592027; HQ591745; HQ591911; HQ592153; HQ592086; EF490247; HQ591842. *Viburnum maculatum* Pant.: K. Schmandt 1294-83B (A). JX049450; JX049455; ; JX049516; JQ805469; JQ805474; JX049489; JQ805300; JQ805554; JX049503. *Viburnum mongolicum* Rehder: M.J. Donoghue s.n. (YU). EF462985; HQ591607; HQ591676; HQ592029; HQ591748; HQ591914; HQ592155; HQ592087; EF490248; HQ591844. *Viburnum rhytidophyllum* Hemsl. ex Forb. & Hemsl.: M.J. Donoghue & R.C. Winkworth 8 (YU). AY265146; HQ591618; HQ591685; HQ592036; HQ591759; HQ591925; HQ592166; HQ592092; AY265192; HQ591850. *Viburnum schensianum* Maxim.: P.W. Sweeney et al. PWS2565 (YU). pending; pending; pending; pending; pending; pending; pending; pending; pending; pending. *Viburnum utile* Hemsl.: P.W. Sweeney et al. PWS2593 (YU). pending; pending; pending; pending; pending; pending; pending; pending; pending; pending. *Viburnum veitchii* C.H. Wright: Bouffourd et al. 27597 (A). HQ591985; HQ591639; HQ591699; HQ592055; HQ591779; HQ591946; -; HQ592106; HQ591817; HQ591861.

***Pseudotinus. Viburnum furcatum*** **Blume ex Hook.f.Thomson: M.J. Donoghue et al. 10 (YU). -; pending; pending; pending; pending; pending; pending; pending; pending; pending.** *Viburnum lantanoides* Michx.: M.J. Donoghue & R.C. Winkworth 2 (YU). AY265135; HQ591596; HQ591668; HQ592020; HQ591737; HQ591903; HQ592146; AY627405; AY265181; EF490279. *Viburnum nervosum* D.Don: P.W. Sweeney et al. PWS2298 (YU). pending; pending; pending; pending; pending; pending; pending; pending; pending; pending. *Viburnum sympodiale* Graebn.: K.F.

Chung KFC-1932 (YU). pending; pending; pending; pending; pending; pending; pending; pending; pending; pending.

***Pluriviburnum.***

***Perplexitinus***

***Urceolata.*** *Viburnum taiwanianum* Hayata: W.-H. Hu et al. 2186 (MO). EF462989; HQ591631; -; HQ592047; HQ591771; HQ591938; HQ592178; HQ592101; EF490253; HQ591855. *Viburnum urceolatum* Siebold & Zucc.: M.J. Donoghue NVI (-). AY265155; HQ591637; HQ591697; HQ592053; HQ591777; HQ591944; -; AY627423; AY265201; HQ591860.

***Amplicrenotinus.*** *Viburnum amplificatum* J. Kern: P.W. Sweeney et al. PWS2149 (YU). KJ795806; KF019753; KF019773; KF019830; KF019795; KF019871; KF019894; KF019850; KF019939; KF019918.

***Solenotinus.*** *Viburnum awabuki* K.Koch: S.-M. Liu et al. 141 (A). HQ591951; HQ591560; -; HQ591990; HQ591704; HQ591867; HQ592111; HQ592060; HQ591783; -. *Viburnum brachybotryum* Hemsl. Et F.B.Forbes & Hemsl.: P.W. Sweeney et al. PWS2222 (YU). pending; pending; pending; pending; pending; pending; pending; pending; -; pending. ***Viburnum brevitubum*** **(P.S.Hsu) P.S.Hsu: P.W. Sweeney et al. PWS2580 (YU). -; pending; pending; pending; pending; pending; pending; pending; pending; pending.** ***Viburnum burmanicum*** **(Rehder) C.Y. Wu: P.W. Sweeney et al. PWS2290 (YU). pending; pending; pending; pending; pending; pending; pending; pending; pending; pending.** *Viburnum cf.*

*chingii* P.S. Hsu: P.W. Sweeney et al. PWS2247 (YU). pending; pending; pending; pending; pending; pending; pending; pending; pending; pending. *Viburnum corymbiflorum* P.S. Hsu & S.C. Hsu: X. Gao 1706 (A). HQ591961; HQ591573; -; -; -; HQ591882; HQ592126; HQ592072; -; -. *Viburnum erubescens* Wall.: Boufford et al. 27190 (A). AY265127; HQ591581; HQ591655; HQ592006; HQ591724; HQ591889; HQ592133; AY627397; AY265173; HQ591831. *Viburnum farreri* Stearn: M.J. Donoghue & R.C. Winkworth 18 (YU). AY265128; HQ591582; HQ591656; HQ502007; HQ591725; HQ591890; HQ592134; AY627398; AY265174; EF490274. *Viburnum foetens* Decne.: M.J. Donoghue WC270 (YU, WU). KF019813; KF019754; KF019774; KF019831; KF019796; KF019872; KF019895; KF019851; KF019940; KF019919. *Viburnum grandiflorum* Wall. ex DC: M.J. Donoghue WC271 (YU). KF019814; KF019755; KF019775; KF019832; KF019797; KF019873; KF019896; KF019852; KF019941; KF019920. *Viburnum henryi* Hemsl.: M.J. Donoghue WC272 (YU). KF019815; KF019756; KF019776; ; KF019798; KF019874; KF019897; KF019853; KF019942; KF019921. *Viburnum odoratissimum* Ker-Gawl.: R. Olmstead 118 (WTU). AY265141; HQ591609; HQ591678; -; HQ591750; HQ591916; HQ592157; AY627411; AY265187; HQ591845. *Viburnum oliganthum* Batalin: D.E. Bouffourd et al. 27175 (A). HQ591971; HQ591610; -; -; HQ591751; HQ591917; HQ592158; HQ592088; HQ591804; HQ591846. *Viburnum sieboldii* Miq.: M.J. Donoghue et al. 7 (YU). pending; pending; pending; pending; pending; pending; pending;

pending; pending; pending. ***Viburnum subalpinum* Hand.-Mazz.: P.W. Sweeney et al. PWS2292 (YU). pending; pending; pending; pending; pending; pending; pending; pending; pending; pending.** *Viburnum suspensum* Lindl.: M.J. Donoghue & R.C. Winkworth 36 (YU). AY265151; HQ591629; HQ591692; HQ592045; HQ591769; HQ591936; HQ592176; AY627419; AY265197; HQ591854. *Viburnum taitoense* Hayata: M.J. Donoghue & K-F. Chung KFC1941 (YU). KF019816; KF019757; KF019777; KF019833; KF019799; KF019875; KF019898; KF019854; KF019943; KF019922. ***Viburnum cf. yunnanense* Rehder: P.W. Sweeney et al. PWS2275 (YU). pending; pending; pending; pending; pending; pending; pending; pending; pending; pending.**

***Lutescentia.*** *Viburnum colebrookeanum* Wall. ex DC: Parker 3220 (A). HQ591959; HQ591570; -; HQ592000; HQ591715; HQ591879; HQ592123; HQ592070; HQ591791; -. *Viburnum hanceanum* Maxim.: P.W. Sweeney et al. PWS2195 (YU). pending; pending; pending; pending; pending; pending; pending; pending; pending; pending. *Viburnum lutescens* Bl.: P.W. Sweeney et al. PWS2077 (YU). pending; pending; pending; pending; pending; pending; pending; -; pending; pending. ***Viburnum pyramidatum* Rehder: P.W. Sweeney et al. PWS2272 (YU). pending; -; pending; pending; pending; pending; pending; pending; pending; pending.**

***Tomentosa.*** *Viburnum plicatum* Thunb.: M.J. Donoghue & R.C. Winkworth 10 (YU). AY265143; HQ591613; HQ591681; HQ592032;

HQ591754; HQ591920; HQ592161; AY627412; AY265189; EF490285.

***Nectarotinus.***

**Tinus.** *Viburnum atrocyaneum* C.B. Clarke: Boufford et al. 34956 (A). HQ591950; HQ591559; HQ591642; HQ591989; HQ591703; HQ591866; HQ592110; HQ592059; HQ591782; HQ591820. *Viburnum calvum* Rehder: H. Li & V. Soukup 934 (A). HQ591955; HQ591565; HQ591644; HQ591995; HQ591709; HQ591872; HQ592116; HQ592066; HQ591788; JX049508. *Viburnum cinnamomifolium* Rehder: P.W. Sweeney et al. PWS2255 (YU). pending; pending; pending; pending; pending; pending; pending; pending; pending; pending. *Viburnum davidii* Franch.: M.J. Donoghue WC269 (YU). KF019821; KF019765; KF019785; KF019841; KF019807; KF019883; KF019906; KF019862; KF019951; KF019930. *Viburnum rigidum* Vent.: W.T. Stearn 1116 (A). HQ591974; HQ591619; HQ591686; HQ592037; HQ591760; HQ591926; -; HQ592093; HQ591807; -. *Viburnum tinus* L.: M.J. Donoghue & R.C. Winkworth 35 (YU). AY265152; HQ591633; HQ591693; HQ592049; HQ591773; HQ591940; HQ592180; AY627420; AY265198; HQ591857. *Viburnum treleasei* Gand.. EF445121; -; -; -; HM850455; -; -; -; EF445122; -.

***Imbricotinus***

***Laminotinus***

***Sambucina.*** *Viburnum glaberrimum* Merr.: P.W. Sweeney et al. PWS2322 (YU). pending; pending; pending; pending; pending;

pending; pending; pending; pending; pending. *Viburnum hispidulum* J. Kern: P.W. Sweeney et al. PWS2136 (YU). -; KF019749; KF019769; KF019826; KF019791; KF019867; KF019890; KF019846; KF019935; KF019914. *Viburnum inopinatum* Craib.: P.W. Sweeney et al. PWS2091 (YU). KJ795808; KF019750; KF019770; KF019827; KF019792; KF019868; KF019891; KF019847; KF019936; KF019915. *Viburnum leiocarpum* P.S. Hsu: P.W. Sweeney et al. PWS2265 (YU). pending; pending; pending; pending; pending; pending; pending; pending; pending; pending. *Viburnum sambucinum* Reinw. ex Blume: P.W. Sweeney et al. PWS2100 (YU). KF019811; KF019751; KF019771; KF019828; KF019793; KF019869; KF019892; KF019848; KF019937; KF019916. ***Viburnum cf. ternatum*** **Rehder: P.W. Sweeney et al. PWS2264 (YU). pending; pending; pending; pending; pending; pending; pending; pending; pending; pending.** *Viburnum vernicosum* Gibbs: P.W. Sweeney et al. PWS2123 (YU). KF019812; KF019752; KF019772; KF019829; KF019794; KF019870; KF019893; KF019849; KF019938; KF019917.

***Corisuccotinus***

***Coriacea.*** *Viburnum beccarii* Gamble: P.W. Sweeney et al. PWS2106 (YU). KF019808; KF019743; KF019766; KF019822; KF019786; KF019863; KF019884; KF019842; KF019931; KF019907. *Viburnum coriaceum* Bl.: P.W. Sweeney et al. PWS2088 (YU). KP281840; KP281810; KP281828; KP281864; KP281818;

KP281854; KP281876; KP281845; KP281893; KP281902.

*Viburnum cylindricum* Buch.-Ham. ex D.Don: P.W. Sweeney et al. PWS2233 (YU). pending; pending; pending; pending; pending; pending; pending; pending; pending; pending. *Viburnum hebanthum* Wight & Arn.: J. Klackenberg 32 (NY). -; HQ591587; HQ591660; HQ592012; HQ591729; HQ591895; HQ592138; HQ592076; HQ591795; HQ591833.

***Succotinus*.** *Viburnum adenophorum* W.W. Smith: Boufford & Barholomew 24402 (A). HQ591948; HQ591558; -; HQ591988; HQ591702; HQ591864; HQ592109; HQ592057; HQ591781; pending. *Viburnum annamensis* N.Fukuoka: P.W. Sweeney et al. PWS2094 (YU). KJ795807; KF019758; KF019778; KF019834; KF019800; KF019876; KF019899; KF019855; KF019944; KF019923. ***Viburnum betulifolium* Batalin: M.J. Donoghue & K-F. Chung KFC1950 (YU). pending; pending; pending; pending; pending; pending; pending; pending; pending; pending. *Viburnum brachyandrum* Nakai: M.J. Donoghue et al. 9 (YU). pending; pending; pending; pending; pending; pending; pending; pending; pending; pending.** *Viburnum brevipes* Rehder: Zhu Dahai et al 110 (MO). -; JQ805283; -; -; JQ805454; JQ805535; -; JQ805366; JQ805608; -. ***Viburnum corylifolium* Hook.f. & Thomson: P.W. Sweeney et al. PWS2249 (YU). -; pending; pending; pending; pending; pending; pending;**

**pending; pending; pending.** *Viburnum dilatatum* Thunb.: P.W. Sweeney et al. PWS2209 (YU). -; pending; pending; pending; pending; pending; pending; pending; -; pending. *Viburnum erosum* Thunb.: M.J. Donoghue et al. 4 (YU). -; pending; pending; pending; pending; pending; pending; pending; pending; pending. ***Viburnum fansipanense*** **J.M.H.Shaw, Wynn-Jones & V.D.Nguyen: P.W. Sweeney et al. PWS2243 (YU). pending; pending; pending; pending; pending; pending; pending; pending; pending; pending.** *Viburnum flavescens* W.W. Smith: Boufford et al. 32758 (A). HQ591962; HQ591583; HQ591657; HQ592008; HQ591726; HQ591891; -; HQ592074; HQ591794; JX049505. ***Viburnum foetidum*** **Wall.: P.W. Sweeney et al. PWS2250 (YU). pending; pending; pending; pending; pending; pending; pending; pending; pending; pending.** ***Viburnum cf. fordiae*** **Hance: P.W. Sweeney et al. 2201 (extracted). pending; -; pending; -; pending; pending; pending; pending; pending; pending.** *Viburnum formosanum* Hayata: M.J. Donoghue & J.M. Hu J-M Hu 2007 (YU). -; KF019760; KF019780; KF019836; KF019802; KF019878; KF019901; KF019857; KF019946; KF019925. *Viburnum hupehense* Rehder: Bartholomew et al. 1286 (A). HQ591964; HQ591588; HQ591661; HQ592013; HQ591730; HQ591896; HQ592139; HQ592077; HQ591796; HQ591834. *Viburnum ichangense* Rehder: Bartholomew et al. 1889 (A).

HQ591965; HQ591589; HQ591662; HQ592014; HQ591731; HQ591897; HQ592140; HQ592078; HQ591797; HQ591835. *Viburnum integrifolium* Hayata: M.J. Donoghue & K-F. Chung KFC1946 (YU). -; KF019761; KF019781; KF019837; KF019803; KF019879; KF019902; KF019858; KF019947; KF019926. *Viburnum japonicum* Spreng: NVI (YU). AY265131; HQ591592; HQ591664; HQ592016; HQ591733; HQ591899; HQ592143; AY627401; AY265177; HQ591837. *Viburnum lancifolium* P.S.Hsu: Zou Huanning (MO). -; -; -; -; JQ805460; JQ805540; -; JQ805373; JQ805612; -. *Viburnum lobophyllum* Graebn.: M.J. Donoghue & R.C. Winkworth 25 (YU). AY265137; HQ591600; HQ591671; HQ592023; HQ591741; HQ591907; HQ592149; AY627407; AY265183; HQ591840. *Viburnum luzonicum* Rolfe: P.W. Sweeney et al. PWS2321 (YU). -; pending; pending; pending; pending; pending; pending; pending; -; pending. ***Viburnum melanocarpum* P.S.Hsu: P.W. Sweeney et al. PWS2560 (YU). pending; pending; pending; pending; pending; pending; pending; pending; pending; pending.** *Viburnum mullaha* Buch.-Ham. ex D.Don: M.J. Donoghue WC274 (YU). KF019819; KF019762; KF019782; KF019838; KF019804; KF019880; KF019903; KF019859; KF019948; KF019927. *Viburnum parvifolium* Hayata: M.J. Donoghue, KFC KFC1953 (YU). KF019820; KF019763; KF019783; KF019839; KF019805; KF019881; KF019904; KF019860;

KF019949; KF019928. *Viburnum phlebotrichum* Siebold & Zucc.: M.J. Donoghue et al. 3 (YU). pending; pending; pending; pending; pending; pending; pending; pending; pending; pending. *Viburnum propinquum* Hemsl.: P.W. Sweeney et al. PWS2188 (YU). pending; pending; pending; pending; pending; pending; pending; pending; pending; pending. *Viburnum sempervirens* K.Koch: P.W. Sweeney et al. PWS2191 (YU). pending; pending; pending; pending; pending; pending; pending; pending; pending; pending. *Viburnum setigerum* Hance: P.W. Sweeney et al. PWS2554 (YU). pending; pending; pending; pending; pending; pending; pending; pending; pending; pending. *Viburnum tashiroi* Nakai: M.J. Donoghue s.n. (YU). -; KF019764; KF019784; KF019840; KF019806; KF019882; KF019905; KF019861; KF019950; KF019929. *Viburnum wrightii* Miq.: M.J. Donoghue et al. 1 (YU). pending; pending; pending; pending; pending; pending; pending; pending; pending; pending.

***Opulus***. *Viburnum edule* (Michx.) Raf.: NVI (-). AY265123; HQ591577; -; -; HQ591720; -; -; AY627393; AY265169; EF490271. *Viburnum koreanum* Nakai: H. Yamaji 5170 (MO). EF462983; -; -; -; -; -; -; HQ592081; EF490246; EF490277. *Viburnum opulus* L.: W.L. Clement 250 (YU). HQ591972; HQ591611; HQ591679; -; HQ591752; HQ591918; HQ592159; -; HQ591805; HQ591847. *Viburnum sargentii* Koehne: M.J. Donoghue & R.C. Winkworth 17 (YU). AY265148; HQ591621; HQ591688; HQ592039; HQ591762; HQ591928; HQ592168; AY627416; AY265194; EF490288.

*Viburnum trilobum* Marshall: Arnold Arboretum 22900A, 0174487 (AA). HQ591983; HQ591635; HQ591695; HQ592051; HQ591775; HQ591942; HQ592182; HQ592104; HQ591815; EF490290.

***Porphyrotinus***

***Mollotinus.*** *Viburnum bracteatum* Rehder: Arnold Arboretum 1067-87A, 0227564 (A). -; HQ591564; HQ591643; HQ591994; HQ591708; HQ591871; HQ592115; HQ592065; KF019933; HQ591822. *Viburnum ellipticum* Hook.: M.J. Donoghue NVI (-). AY265125; HQ591579; HQ591653; HQ592004; HQ591722; -; HQ592131; AY627395; AY265171; HQ591830. *Viburnum molle* Michx.: M.J. Donoghue & R.C. Winkworth 5 (YU). AY265139; HQ591606; HQ591675; -; HQ591747; HQ591913; HQ592154; AY627409 ; AY265185; EF490281. *Viburnum rafinesquianum* Schult.: M.J. Donoghue & R.C. Winkworth 4 (YU). AY265145; HQ591617; HQ591684; HQ592035; HQ591758; HQ591924; HQ592165; AY627414; AY265191; HQ591849.

***Oreinodentinus***

***Dentata.*** *Viburnum dentatum* L.: M.J. Donoghue & R.C. Winkworth 33 (YU). AY265121; HQ591574; HQ591651; HQ592002; HQ591718; HQ591884; HQ592128; AY627391; AY265167; HQ591827. *Viburnum recognitum* Fernald: Arnold Arboretum 1471-83B, 00192902 (A). JQ805189; JQ805261; JX049465; KF019824; JQ805387; JQ805507; JX049490; JQ805337;

JQ805585; JX049504. *Viburnum scabrellum* Chapman: M.J. Donoghue 82 (YU). JQ805190; JQ805262; KP281836; KP281871; KP281824; JQ805508; KP281887; JQ805338; JQ805586; KP281910.

***Oreintotinus. Viburnum acutifolium*** **Benth.: P.W. Sweeney et al. PWS3059 (YU). pending; pending; pending; pending; pending; pending; pending; pending; -; pending.** *Viburnum anabaptista* Graebn.: P.W. Sweeney et al. PWS2160 (YU, ANDES). -; KP281812; KP281829; KP281866; KP281820; KP281856; KP281878; KP281847; KP281895; KP281904. *Viburnum australe* Morton: M.A. Carranza et al. 2064 (MO). JQ805157; JQ805235; -; -; JQ805393; -; KP281879; JQ805304; KP281896; -. *Viburnum ayavacense* Kunth: E. Fernandez et al. 1930 (MO). JQ805162; JQ805238; KP281830; -; JQ805399; KP281857; KP281880; JQ805309; JQ805561; -. *Viburnum blandum* C.V. Morton: M.J. Donoghue 464 (YU). HQ591952; HQ591562; -; HQ591992; HQ591706; HQ591869; HQ592113; HQ592062; HQ591785; -. *Viburnum caudatum* Greenm.: M.J. Donoghue 64 (YU). HQ591957; -; -; -; -; HQ591875; HQ592119; HQ592068; HQ591790; HQ591825. *Viburnum ciliatum* Greenm.: M.J. Donoghue 48 (YU). -; JQ805240; pending; pending; JQ805401; pending; pending; JQ805311; JQ805563; -. *Viburnum costaricanum* Hemsl.: M.J. Donoghue 85 (YU). JQ805164; -; KP281831; -; -; JQ805482; -; -; JQ805564; KF019909. *Viburnum* *discolor* Benth.: M. Veliz, N. Gallardo, M. Vasquez 35-99 (MO). JQ805166; JQ805241; -; -; JQ805402; JQ805485; KF019886; JQ805314; pending; -. *Viburnum disjunctum* C.V. Morton: M.J. Donoghue 700 (YU). KF019810; KF019745; -; -; KF019788; -; KF019887; KF019844; -; KF019910. *Viburnum divaricatum* Benth.: P.W. Sweeney et al. PWS1773 (YU, QCNE). JQ805167; JQ805242; KP281832; KP281867; JQ805404; JQ805487; KP281882; JQ805317; JQ805567; KP281905. ***Viburnum fuscum* Hemsl.: P.W. Sweeney et al. PWS3054 (YU). pending; pending; pending; pending; pending; pending; pending; -; pending; pending.** *Viburnum glabratum* Kunth: P.W. Sweeney et al. 2152 (YU, ANDES). KP281841; KP281813; KP281833; KP281868; KP281821; KP281858; KP281883; KP281848; KP281897; KP281906. *Viburnum hallii* Killip & A.C. Smith: P.W. Sweeney et al. PWS1626 (YU, QCNE). JQ805173; JQ805248; -; pending; JQ805410; JQ805492; pending; JQ805322; JQ805572; pending. *Viburnum hartwegii* Benth.: M.J. Donoghue 40 (YU). AY265130; HQ591586; HQ591659; HQ592011; -; HQ591894; HQ592137; AY627400; AY265176; HQ591832. *Viburnum jamesonii* Killip & A.C. Smith: P.W. Sweeney et al. 1636 (YU). HQ591966; HQ591591; HQ591663; HQ592015; HQ591732; HQ591898; HQ592142; HQ592080; HQ591798; HQ591836. *Viburnum jelskii* Zahlbr.: Ynes Mexia 8224 (GH). pending; pending; pending; pending; pending; pending;

pending; -; pending; pending. *Viburnum jucundum* C.V. Morton: M.J. Donoghue 244 (YU). AY265132; HQ591593; HQ591665; HQ592017; HQ591734; HQ591900; -; AY627402; AY265178; HQ591838. *Viburnum lasiophyllum* Benth.: P.W. Sweeney et al. 2174 (YU, ANDES). -; KP281814; KP281834; KP281869; KP281822; KP281859; KP281884; KP281849; -; KP281907. *Viburnum lautum* C.V. Morton: M.J. Donoghue 72 (YU). HQ591967; HQ591597; HQ591669; HQ592021; HQ591738; HQ591904; HQ592147; HQ592082; HQ591799; HQ591839. *Viburnum loeseneri* Graebn.: M.J. Donoghue 2547 (YU). HQ591968; HQ591601; -; HQ592024; HQ591742; HQ591908; HQ592150; HQ592084; HQ591801; -. *Viburnum microcarpum* Schlecht. & Cham.: F. Ventura A. 819 (NY). JQ805178; -; -; -; JQ805414; JQ805496; -; JQ805327; KP281898; -. *Viburnum microphyllum* Hemsl.: M.J. Donoghue 2492 (YU). -; pending; -; -; KP281823; KP281860; KP281885; KP281850; -; KP281908. *Viburnum obtectum* J.H.Vargas: P.W. Sweeney et al. PWS1701 (YU, QCNE). JQ805179; JQ805252; -; -; JQ805415; JQ805497; -; JQ805328; JQ805576; -. *Viburnum obtusatum* D.N.Gibson: P.W. Sweeney et al. PWS3100 (YU). pending; pending; pending; pending; pending; pending; pending; pending; -; pending. *Viburnum pastasanum* Diels: P.W. Sweeney et al. PWS1799 (YU, QCNE). HQ591982; HQ591634; HQ591694; HQ592050; HQ591774; HQ591941;

HQ592181; HQ592103; HQ591814; HQ591858. *Viburnum pichinchense* Benth.: P.W. Sweeney et al. PWS1669 (YU, QCNE). JQ805184; JQ805257; KP281835; KP281870; JQ805420; JQ805502; KP281886; JQ805332; JQ805580; -. *Viburnum seemenii* Graebn.: M. Lewis 37409 (GH). JQ805193; JQ805263; -; -; JQ805426; JQ805509; -; JQ805340; JQ805587; -. *Viburnum stellato-tomentosum* Hemsl.: M.J. Donoghue MJD640 (YU). -; KF019747; -; -; KF019789; KF019865; KF019888; KF019845; -; KF019911. *Viburnum stenocalyx* Hemsl.: M.J. Donoghue 60 (YU). HQ591978; HQ591626; -; HQ592043; HQ591767; HQ591933; HQ592173; HQ592097; HQ591810; KF019912. *Viburnum stipitatum* J.H.Vargas: P.W. Sweeney et al. PWS1708 (YU, QCNE). JQ805195; JQ805265; -; -; JQ805428; JQ805512; -; JQ805343; JQ805589; -. *Viburnum subsessile* Killip & A.C. Smith: P.W. Sweeney et al. 2172 (YU, ANDES). KP281842; KP281815; KP281837; KP281872; KP281825; KP281861; KP281888; KP281851; -; KP281911. ***Viburnum sulcatum* Hemsl.: P.W. Sweeney et al. PWS3053 (YU). pending; pending; pending; pending; pending; pending; pending; pending; -; pending.** *Viburnum tinoides* L.: P.W. Sweeney et al. 2167 (YU, ANDES). KP281843; KP281816; KP281838; KP281873; KP281826; KP281862; KP281889; KP281852; KP281899; KP281912. *Viburnum triphyllum* Benth.: P.W. Sweeney et al. PWS1783 (YU, QCNE). HQ591984; HQ591636;

HQ591696; HQ592052; HQ591776; HQ591943; HQ592183; HQ592105; HQ591816; HQ591859. *Viburnum undulatum* Killip & A.C. Smith: P.W. Sweeney et al. 2164 (YU, ANDES). KP281844; KP281817; KP281839; KP281874; KP281827; KP281863; KP281890; KP281853; KP281900; KP281913. *Viburnum venustum* C.V.Morton: W. Haber & W. Zuchowski 11072 (MO). JQ805206; JQ805278; -; ; JQ805441; JQ805525; KP281891; JQ805355; KP281901; KP281914. *Viburnum villosum* Sw.: M.J. Donoghue 628 (YU). -; JQ805280; -; KP281875; JQ805443; JQ805527; KP281892; JQ805357; JQ805600; -.

**Supplement 3. Placement of unsequenced species based on morphological traits**

We have been unable to obtain DNA sequences for 10 extant species. Aside from *V. hondurense* (where we have tried and failed to obtain DNA extractions of sufficient quality), these species are very poorly known in the wild and in herbaria, and are only tentatively retained in our world-wide monographic treatment of *Viburnum* (Donoghue et al., in prep.). The phylogenetic placements of these 10 species were variously constrained based on morphological characters as described below with reference to Figs. S6. Note that we have used apomorphies for the placement of species within a major clade (positive constraints), and in some cases we have used additional characters to exclude their placement within one or more strongly supported sub-clades (negative constraints).

*Viburnum tengyuehense, V. shweilense and V. wardii* are placed in the *Solenotinus* clade based their panicle inflorescences (with just two opposite rays at the first inflorescence node; Clement et al., 2014; Fig. S6a) Within *Solenotinus* they are constrained outside of the *V. suspensum*-*V. taitoense* clade based on their lack of thick evergreen leaves with crenulate (*V. suspensum* of the Ryuku Islands) or sharp-pointed (*V. taitoense* of Taiwan) teeth. They are also constrained outside of the *V. farreri*-*V. foetens*-*V. grandiflorum* clade. The latter three species are characterized by apomorphies including tubular corollas with the stamens inserted at different heights above the middle of the corolla tube, and flowers that open before the leaves expand.

*Viburnum garretii* and *V. junghunii* are placed in the *Lutescentia* clade based on shared branching architecture (Fig. S6a). All members of this clade produce monopodial plagiotropic axes, with inflorescences characteristically borne ("double-file", i.e., on both sides of the main axis) on shortened lateral branches (Donoghue, 1981; Edwards et al.,

2014). These species also are characterized by endocarps with a broad ventral groove (Jacobs et al., 2008). Within *Lutescentia*, *V. garretii* and *V. junghunii* are excluded from the *V. plicatum*-*V. hanceanum* clade. These latter two species bear enlarged and asymmetrical sterile flowers around the periphery of their inflorescences (Clement et al., Park and Donoghue, in press); sterile marginal flowers are lacking in all other species of *Lutescentia*.

*Viburnum chunii*, *V. dalzielii*, *V. hainanense*, and *V. longiradiatum* are constrained to the *Succotinus* clade based on their fruits, which are red and juicy at maturity, with flattened endocarps with shallow grooves on both the dorsal and ventral surfaces (Jacobs et al., 2008; Fig. S6a). They also connect with this clade (and several smaller related clades) based on the production of extrafloral nectaries within the leaf blade (laminar EFNs: Webber et al., 2012; Clement et al., 2014). We allowed these four species to connect anywhere within *Succotinus* (i.e., we know of no basis for excluding them from any subclades).

*Viburnum hondurense* belongs to the Neotropical *Oreinotinus* clade based primarily on its fruits (Fig. S6d). These mature simultaneously to a dark purple/black color, have a relatively thin and mealy mesocarp, and a more-or-less round endocarp in cross section with only a very shallow ventral groove (Jacobs et al., 2008). This placement is supported also by the presence of one or more pairs of marginal extrafloral nectaries at the base of the leaf, which characterizes the more inclusive *Porphyrotinus* clade. For purposes of the present analyses, we excluded *V. hondurense* from the South American clade that has been well supported in prior phylogenetic analyses. This decision reflects our desire to evaluate the monophyly of the South American species based on sequenced species, independent of the inclusion of a rogue unsequenced species. Obtaining DNA from *V. hondurense* is a high priority, especially to more critically evaluate the placement of the Jamaican/Cuban species.

*References*

Clement, W., M. Arikake, P. Sweeney, E. J. Edwards, and M. J. Donoghue. 2014. A chloroplast tree for *Viburnum* (Adoxaceae) and its implications for phylogenetic classification and character evolution. Amer. J. Bot. 101: 1029-1049.

Donoghue, M.J. 1981. Growth patterns in woody plants with examples from the genus *Viburnum*. Arnoldia 41:2-23.

Edwards, E. J., D. Chatelet, L. Sack, and M. J. Donoghue. 2014. Leaf life span and the leaf economic spectrum in the context of whole plant architecture. Jour. Ecology 102: 328-336.

Jacobs, B., M. J. Donoghue, F. Bouman, S. Huysmans, and E. Smets. 2008. Evolution and phylogenetic importance of endocarp and seed characters in *Viburnum* (Adoxaceae). Int. J. Plant Sci. 169: 409-431.

Park, B. and M. J. Donoghue. Phylogenomic insights into the independent origins of sterile marginal flowers in *Viburnum* (Adoxaceae). Amer. J. Botany (in review).

Weber, M. G., M. J. Donoghue, W. L. Clement, and A. A. Agarwal. 2012. Phylogenetic and experimental tests of interactions among mutualistic plant defense traits in *Viburnum* (Adoxaceae). Amer. Nat. 180: 450-463.

**Supplement 4. Fossil pollen morphology and biome assignments**

Fossil leaves identified as *Viburnum* species from the Paleocene and Eocene were reassigned to other plant clades in a series of studies by Manchester and colleagues (Manchester et al., 1999; Manchester, 2002a, b; Manchester and Hickey, 2007). Although it remains possible that some of these specimens truly are viburnums, this requires further analysis, and at this stage it is prudent to disregard them. Instead, we have incorporated five fossil pollen grains for dating purposes, scoring their exine morphologies, their geographic localities, and their likely biome affinities. These grains are assigned to one of the three *Viburnum* exine states recognized by Donoghue (1985; also see Bohnke-Gutlein and Weberling, 1981); these types (Ia, Ib, and Ic) are described in the text and illustrated in Figs. S7 and S8. When SEMs of the fossil grains have been published (Iceland, Colorado, British Columbia,), the assignment to one of these states is straightforward. Light micrographs (Paris, Northwest Territories) are more difficult to interpret, but permit assignments with somewhat less confidence.

Our biome state assignments are based on analyses of the collecting localities and their inferred climates. In part, this relies on assessing associated element of the paleoflora and the climatic preferences of their closest living relatives (assuming niche conservatism). This can be difficult in view of the fact that fossils collected from a site often include elements from a set of adjacent environments in what may have been a complex landscape. Nevertheless, we rely on assessments made in the literature and on our own judgements about these paleofloras and their implications for climate and biome occupancy.

**Iceland (~15 MYA).** Based on Plate 4.9 in Denk et al. (2011) we scored these grains as type Ic of Donoghue (1985). This state evolved from Ib within the *Lentago* clade, and independently from Ib within the *Euviburnum* clade (Donoghue, 1985; Clement et al., 2014), and, as described in the text, we imposed an either-or-constraint to assure the placement of these grains in one of those two clades.

Denk et al. (2011) described a "diverse humid warm temperate flora" in Iceland between 12 and 15 Ma, and they argued that Iceland may have served at that time as a segment of a corridor for plant movement between Europe and Eastern North America. Several elements of the paleoflora do suggest a warmer climate, such as *Cercidophyllum*, *Sciadopytis*, *Magnolia*, *Pterocarya*, *Tetracentron*, *Liquidambar*, *Calycanthaceae*, and several Lauraceae. However, the species list for that time period includes mainly plants that we associate with cold temperate forests today. *Therefore, we scored this fossil as having occupied either a warm temperate (W) or a cold temperate (F) forest.*

**Florissant (Colorado, USA, ~34 MYA).** Based on Plate 22 G-L in Bouchal (2013), we scored fossil *Viburnum* pollen from the Florissant formation as type Ia of Donoghue (1985). As this is the presumed ancestral exine morphology for *Viburnum* (Clement et al., 2014), we placed no constraints on its phylogenetic position.

Based on Leopold et al. (2007), and especially Zaborac-Reed and Leopold (2016), *we scored the biome of these fossils as warm temperate (W).* These authors described the presence of many elements of modern warm temperate forests, including Lauraceae, Arecaceae (palms), Meliaceae, Podocarpaceae, Sapindaceae, Sterculiaceae, and *Torreya*. They calculated that more than 55% of the plants were "warm loving", many with their closest living relatives in Southern China or in the humid forests of Mexico. Their analyses estimated a mean annual temperature (MAT) of between 14-18 C for the site, which is squarely in the mesophyllous range and likely supported a broad-leaved warm temperate forest (with conifers also present). Although it was initially thought to represent a higher elevation flora, Zaborac-Reed and Leopold (2016) concluded that it was likely lower elevation (<1.5 km).

**Axel Heiburg Island (Northwest Territories, Canada, ~45 MYA).** Based on Plate 3 Figs. 19-21 in McIntyre (1991) we scored these grains as Type Ib of Donoghue (1985). On this basis we constrained its placement to the *Valvatotinus* clade. McIntyre (1991) compared these grains to *V. cassinoides* of the *Lentago* clade, but this exine type is widespread and ancestral in *Valvatotinus*.

Based on McIntyre (1991), Jahren (2007), and Greenwood et al. (2010) *we coded the biome of these fossils as warm temperate (W).* Based on leaf morphological and nearest relative analyses, these and other authors have supported the view that the MAT at that time was above 12 C, and most likely between 13-18 C, in the mesothermal range. Hard and prolonged freezes were therefore likely rare, despite the dark winters at this high latitude. Based on leaf morphology and nearest relative analyses, Greenwood et al (2010) estimated cold month mean temperatures of ~4 C, and compared Eocene Axel Heiburg with modern coastal conditions in the Pacific Northwest.

**Paris Basin (France, ~45 MYA).** Based on Plate XIV Figs. 16-19 in Gruas-Cavagnetto (1978) we scored this fossil pollen as 1b and constrained it to *Valvatotinus*. Though it is difficult without a scanning electron micrograph, the light micrograph indicates the presence of the continuous but scabrate muri characteristic of this exine type. Gruas-Cavagnetto (1978) compared it to *V. punctatum*, a subtropical Asian species of *Valvatotinus* with Ib exine.

Huyghe et al. (2015) estimated (based mainly on fossil marine animals) that sea surface temperatures were >20C in the Paris Basin during this interval in the mid-Eocene, with a drop in the Late Eocene to perhaps 12 C. On the basis of this mesothermal range, *we infer that the biome was warm temperate (W).* Based on an analysis of the fossil pollen evidence, Gruas-Cavagnetto (1978) concurred that the climate was hot in the Paris Basin in the lower and middle Eocene and likely supported a warm temperate or subtropical forest that included a variety of temperate trees along with tropical elements such as Marratiaceae, Arecaceae, Bombacaceae, and Theaceae.

**Princeton Chert (Alendy, British Columbia, Canada, 47 MYA).** Based on Fig. 20 in Manchester et al. (2015) we score these grains as the ancestral 1a type of Donoghue (1985), and, accordingly, placed no morphological constraints on its phylogenetic position.

Greenwood et al. (2005) estimated MAT's between 13-15 C across six sites in the southern part of the Okanagan Highlands of British Columbia and adjacent Washington, and cold month mean temperatures from 3.5-5.8 C. Noting the presence of palms (Arecaceae) at these sites in British Columbia, Moss et al. (2005) inferred a coldest month mean temperature of >5 C. These studies suggest a mesothermal regime, without prolonged freezing, but bordering on microthermal at some sites. Based on these analyses and the presence of Cycadaceae, Arecaceae, Theaceae, Sapotaceae, and Musaceae in these paleofloras at that time, *we scored the biome as warm temperate (W)*.

*References*

Bohnke-Gutlein E. and F. Weberling. 1981. Palynologische Untersuchungen an Caprifoliaceae. I. Sambuceae, Viburneae und Diervilleae. Tropische und Subtropische Pflanzenwelt 34: 131–189.

Bouchal. J.M. 2013. The microflora of the uppermost Eocene (Preabonian) Florissant Formation, a combined method approach. MS thesis, University of Vienna.

McIntyre, D. J. 1991. Pollen and spore flora of an Eocene forest, eastern Axel Heiberg Island, N.W.T. In R. L. Christie and N. J. McMillan (eds.), Geological Survey of Canada Bulletin 403: 83-97.

Denk, T., F. Grımsson, R. Zetter, and L. A. Sımonarson. 2011. Late Cainozoic Floras of Iceland: 15 Million Years of Vegetation and Climate History in the Northern North Atlantic. Topics Geobiol., Vol. 35. Springer Netherlands, Dordrecht.

Donoghue, M. J. 1985. Pollen diversity and exine evolution in *Viburnum* and the Caprifoliaceae *sensu lato*. Journal of the Arnold Arboretum 66: 421–469.

Greenwood, D. R., S. B. Archibald, R. W. Mathewes, and P. T. Moss. 2005. Fossil biotas from the Okanagan Highlands, southern British Columbia and northeastern Washington State: climates and ecosystems across an Eocene landscape. Canadian Journal of Earth Science 42: 167–185.

Greenwood, D. R., Basinger, J. F., and Smith, R. Y. 2010. How wet was the Arctic Eocene rain forest? Estimates of precipitation from Paleogene Arctic macrofloras. Geology 38: 15-18.

Gruas-Cavagnetto, C. 1978. Étude palynologique de L'Éoècene du Bassin Anglo-Parisien. Mémoires de la Société Géologique de France (Nouvelle Série) 131: 1–64, 15 plates.

Huyghe, D., Lartaud, F., Emmanuel, L., Merle, D., and Renard, M. 2015. Palaeogene climate evolution in the Paris Basin from oxygen stable isotope (δ18O) compositions of marine molluscs. Journal of the Geological Society, 172(5), 576-587.

Jahren, A. H. 2007. The Arctic Forest of the Middle Eocene. Annual Review of Earth and Planetary Sciences 35: 509–540.

Manchester, S. R. 2002. Leaves and fruits of *Davidia* (Cornales) from the Paleocene of North America. Systematic Botany 27: 368–382.

Manchester, S. R, M. A. Akhmetiev, and T. M. Kodrul. 2002. Leaves and fruits of *Celtis aspera* (Newberry) comb. nov. (Celtidaceae) from the Paleocene of North America and Eastern Asia. International Journal of Plant Sciences 163: 725–736.

Manchester, S. R., P. R. Crane, and L. B. Golovneva. 1999. An extinct genus with affinities to extant *Davidia* and *Camptotheca* (Cornales) from the Paleocene of North America and Eastern Asia. International Journal of Plant Sciences 160: 188–207.

Manchester, S. R., F. Grímmson, and Reinhard Zetter. 2015. Assessing the fossil record of Asterids in the context of our current phylogenetic framework. *Annals of the Missouri Botanical Garden* 100: 329-363.

Manchester, S. R. and L. J. Hickey. 2007. Reproductive and vegetative organs of *Browniea* gen. n. (Nyssaceae) from the Paleocene of North America. *International Journal of Plant Sciences* 168: 229–249.

McIver, E. E. and J. F. Basinger. 1999. Early Tertiary floral evolution in the Canadian high Arctic. Annals of the Missouri Botanical Gardens 86: 523–545.

McIntyre, D. J. 1991. Pollen and spore flora of an Eocene forest, eastern Axel Heiberg Island, N.W.T. In: Tertiary Fossil Forests of the Geodetic Hills Axel Heiberg Island, Arctic Archipelago, Christie RL, McMillan NJ, eds, Geological Survey of Canada Bulletin 403: 83–97.

Moss, P. T., Greenwood, D. R., & Archibald, S. B. 2005. Regional and local vegetation community dynamics of the Eocene Okanagan Highlands (British Columbia - Washington State) from palynology. Canadian Journal of Earth Sciences 42: 187–204.

Kalkreuth, W. D., D. J. McIntyre, and R. Richardson. 1993. The geology, petrography and palynology of Tertiary coals from the Eureka Sound Group at Strathcona Fiord and Bache Peninsula, Ellesmere Island, Arctic Canada. Journal of Coal Geology 24: 75–111.

Leopold, E. B. and S. Clay-Poole. 2001. Florissant leaf and pollen floras of Colorado compared: climatic implications. In: Evanoff, E, K. Gregory, K. Wodziski, K. Johnson, eds. Fossil Flora and Stratigraphy of the Florissant Formation, Colorado. Proc. Denver Museum of Nature and Science Series 4 No. 1. Symposium Volume. Denver: Denver Museum of Nature & Science, pp. 17-70.

Zaborac-Reed, S. J. and Leopold, E. B. 2016. Determining the paleoclimate and elevation of the late Eocene Florissant flora: support from the coexistence approach. Canadian Journal of Earth Sciences 53: 565–573.

**Supplement 5. Bayesian phylogenetic analysis**

We used the fossilized birth-death process to model how lineages diversified and deposited fossil occurrences to produce the total of 163 living + 5 fossil taxa. We assigned a lognormal prior to the net diversification rate (birth rate minus death rate, $\lambda - \mu$) centered on the value that generates 163 taxa after 65 Myrs, and with a shape parameter (lognormal standard deviation) of 0.5 (Nee et al., 1994). The relative turnover rate (death rate divided by birth rate, $\mu/\lambda$) was assigned a Beta(2,2) distribution. With just five of 168 taxa being fossil samples, we applied an informative prior to the fossil sampling rate, $\psi = s \times \lambda$ where $s \sim$ Beta(1, 10) such that the prior mean of $\psi$ is $1/(1+10)\times\lambda$. Fossil ages were treated as uniformly distributed random variables bounded by the appropriate age estimates per specimen (topological constraints for fossils are described below). All currently accepted living *Viburnum* species are represented in the phylogeny, so we set the sampling probability $\rho = 1$. Finally, we converted the posterior stem age mean and variance of Adoxaceae from Bell and Donoghue (2005) into a truncated normal distribution ($\mu = 71$, $\sigma = 10.8$, bounded to $52.71 < x < 85.74$), then applied this density as a secondary calibration to the origin time (i.e., the time when the first *Viburnum* lineage originated).

Molecular variation was modeled by a substitution process. Branch clock rate heterogeneity was modeled by a discrete uncorrelated lognormal prior (UCLN; Drummond et al. 2006) with $k = 50$ quantiles and an exponential rate-1 hyperprior on the UCLN standard deviation. The mean value for the relaxed clock was uniform over orders of magnitude to minimize its influence on the dating estimates, Loguniform(1E-6, 1E2). Each partitioned locus evolved under an independent GTR substitution process (Tavaré 1986) with Gamma-distributed site-rate heterogeneity (+Γ4; Yang 1994). Base frequencies and exchangeability rates were assigned flat Dirichlet priors. The shape/scale parameters for the site-rate variation model were assigned flat priors of Uniform(0, 50). Substitution rates were further relaxed across partitioned loci using rate-multipliers, with the first locus having the rate-multiplier of 1, and all remaining loci having independent rate-multipliers, each with the prior Gamma(2, 2).

Species ranges evolved under the Dispersal-Extinction-Cladogenesis (DEC) model of Ree et al. (2005). We classify area pairs as adjacent if they are roughly as close as Russia and Canada are today. Dispersal events between adjacent areas occurred at the relative rate of 1, while dispersal between non-adjacent areas occurred at a fraction of that rate, $0 < z < 1$, where z is an estimated parameter with a Uniform(0,1) prior. Area adjacencies are initially set to SE As—E As, E As—Eur, E As—N Am, Eur—N Am, and N Am—C Am. Two adjacencies change with time (Ree & Smith, 2008, Landis 2017): C Am—S Am become adjacent at 15 Ma (Montes et al., 2012, Bacon et al., 2015) and Eur—N Am lose adjacency at 12 Ma (Denk et al., 2011), both events dated with conservative age estimates to minimize the effect of the events on historical biogeography.

Ranges were constrained to be fewer than three areas in size. All areas shared the same relative extirpation rate, which was assigned a lognormal prior with mean of 1 and standard deviation of 0.1. We multiplied the relative anagenetic rates of dispersal and extirpation by an uninformative biogeographic clock whose prior is uniform over orders of magnitude, Loguniform(1E-6, 1E2). Our biogeographic dataset does not include ranges of size zero, so we condition our anagenetic transition probabilities to account for this acquisition bias (Landis et al., 2018). During cladogenesis, daughter lineages inherit ranges that are either sympatric (within a region) or allopatric (among regions). Cladogenetic range evolution only occurs at speciation events, which are represented in the phylogeny by internal nodes with two daughter lineages; no cladogenesis occurs for internal nodes representing sampled ancestors under the fossilized birth death process. Each cladogenetic pattern is assigned the weight of its type, which are normalized by the sum of weights for the ancestral range to yield the cladogenetic transition probability matrix (Matzke, 2014). Sympatric events are assigned the weight parameter $w_s$ with a Uniform(0,1) prior, while allopatry events have the weight $w_a = 1 - w_s$.

Shifts between biomes were modeled using a four-state continuous-time Markov process with twelve asymmetric rates. We assigned a flat Dirichlet prior to the transition rates and set the root node's stationary frequencies as uniform. Finally, the overall rate of the process was governed by a biome shift clock with a prior that, like the molecular and biogeographic clock priors, is uniform over orders of magnitude, Loguniform(1E-6, 1E3).

We constrained the phylogenetic relationships for three subsets of our taxa, each requiring different justifications and assumptions. Phylogenies that did not obey the clade constraints specified below were assigned zero prior probability, so they were not sampled in the posterior.

*References*

Bacon, C. D., D. Silvestro, C. Jaramillo, B. T. Smith, P. Chakrabarty, and A. Antonelli. 2015. Biological evidence supports an early and complex emergence of the Isthmus of Panama. Proceedings of the National Academy of Sciences, 112: 6110-6115.

Bell, C. D. and M. J. Donoghue. 2005. Dating the Dipsacales: comparing models, genes, and evolutionary implications. Amer. Jour. Bot. 92: 284-296.

Denk, T., F. Grımsson, R. Zetter, and L. A. Sımonarson. 2011. Late Cainozoic Floras of Iceland: 15 Million Years of Vegetation and Climate History in the Northern North Atlantic. Topics Geobiol., Vol. 35. Springer Netherlands, Dordrecht.

Drummond, A. J., Ho, S. Y., Phillips, M. J., & Rambaut, A. 2006. Relaxed phylogenetics and dating with confidence. PLoS Biology 4:e88.

Landis, M. J. 2017. Biogeographic dating of speciation times using paleogeographically informed processes. Systematic Biology, 66: 128-144.

Landis, M. J., W. A. Freyman, B. G. Baldwin. 2018. Retracing the Hawaiian silversword radiation despite phylogenetic, biogeographic, and paleogeographic uncertainty. Evolution 72: 2343-2359.

Montes, C., A. Cardona, R. McFadden, S. E. Morón, C. A. Silva, S. Restrepo-Moreno, D. A. Ramírez, N. Hoyos, J. Wilson, D. Farris, and G. A. Bayona. 2012. Evidence for middle Eocene and younger land emergence in central Panama: implications for Isthmus closure. Geological Society of America Bulletin, 124: 780-799.

Nee, S., May, R. M., & Harvey, P. H. 1994. The reconstructed evolutionary process. *Philos Trans R Soc Lond B Biol Sci* 344: 305-311.

Ree, H. R., B. R. Moore, C. Webb, and M. J. Donoghue. 2005. A likelihood framework for inferring the evolution of geographic range on phylogenetic trees.  Evolution 59: 2299-2311.

Ree, R. H., & Smith, S. A. 2008. Maximum likelihood inference of geographic range evolution by dispersal, local extinction, and cladogenesis. Systematic Biology 57: 4-14.

Tavaré, S. 1986. Some probabilistic and statistical problems in the analysis of DNA sequences. Lectures on Mathematics in the Life Sciences 17: 57-86.

Yang, Z. 1994. Maximum likelihood phylogenetic estimation from DNA sequences with variable rates over sites: approximate methods. Journal of Molecular evolution, 39: 306-314.

**Supplement 6. Extended discussion of phylogenetic and ancestral biome estimates**

*Phylogenetic differences compared with previous cpDNA trees.* Our RAD-seq and combined results differ in six ways from previous analyses based primarily on cpDNA data (Winkworth and Donoghue, 2005; Clement et al., 2011; Clement et al., 2014; Spriggs et al. 2015; Lens et al., 2016). (1) As noted in the text, our trees are equivocal with respect to the placement of *V. clemensiae*. One new possibility is that *V. clemensiae* is an early diverging branch within *Viburnum* (Fig. 2A,B). (2) *Pseudotinus* and *Urceolata* are linked in our analyses, in line with morphological characters (cf. Eaton et al., 2017). (3) Within *Valvatotinus*, *Punctata* connects with *Lentago* in our analyses, whereas it is linked with *Euviburnum* based on cpDNA. Here, too, our result is more consistent with morphological characters (e.g., the production of peltate scales in *Punctata* and *Lentago*; Clement and Donoghue 2011; Clement et al. 2014). (4) *Coriacea* and *Sambucina* form a clade in our analyses, whereas they are paraphyletic based on cpDNA, with *Coriacea* being more closely related to *Lobata*+*Succotinus*. (5) *V. grandiflorum*, *V. foetens*, and *V. farreri* form a clade within *Solenotinus* in our analyses, which is in turn linked with *V. suspensum* and *V. taitoense*, consistent with data on chromosome numbers (2N=16 in this clade; Egolf, 1962). In contrast, *V. grandiflorum* and *V. foetens* are paraphyletic at the base of *Solenotinus* based on cpDNA, and widely separated from *V. farreri*. (6) Our analyses strongly support the monophyly (versus paraphyly in cpDNA analyses) of the *Lobata* clade, which includes *V. kansuense*, *V. orientale*, and *V. acerifolium*, again consistent with morphology (e.g., tri-lobed leaves).

*Phylogenetic differences compared with Eaton et al. 2017.* Compared to the SVDquartets tree presented in Eaton et al. (2017) there are three minor differences. (1) *V. erosum* is linked directly with *V. brachyandrum* (newly added), and these are sister to *V. anamensis* and *V. sempervirens*, whereas in Eaton et al. *V. erosum* appeared as sister to a clade containing six *Succotinus* species. (2) In *Sambucina*, Eaton et al. (2017) found that *V. glaberrimum* and *V. vernicosum* form a clade with *V. cylindricum* and *V. coriaceum*, where we place *V. glaberrimum* and *V. vernicosum* elsewhere within *Sambucina*, as a clade sister to *V. beccarii*. (3) We place *V. sempervirens* and *V. anamensis* as a clade nested deep within *Succotinus*, where in Eaton et al. this clade diverges just after the branching of *V. integrifolium*.

*Ancestral biome estimates in comparison to Lens et al. (2016).* Lens et al. (2016) argued that *Viburnum* originated in cold temperate forests, whereas we conclude that it originated in warm temperate (lucidophyllous) forests. Lens et al. (2016) reached a different conclusion for three reasons. The first concerns sampling and tree topology. They included 97 *Viburnum* species (as compared to our 163), and they based their analysis on a limited (mainly cpDNA) molecular dataset. Although their published tree is broadly similar to ours, it differs in ways that influence the reconstruction of ancestral biome. The presence or absence of particular species plays a key role. For example, absent from their analyses are a number of tropical and warm temperate species that are spread widely in the phylogeny, e.g., *V. amplificatum*, *V. lepidotulum*, *V. brachybotrium*, and the entire *Sambucina* clade of SE Asia.

Second, their climate scorings for the modern species differ in a number of instances from ours. Their use of GBIF locality records yielded erroneous conclusions in several important cases. The reasons differ from case to case, but *V. nudum* is emblematic of one important problem. In this case, there is long-standing taxonomic confusion over whether *V. nudum* and *V. cassinoides* should be recognized as separate species. Consequently, many specimens of the more northern, cold-adapted *V. cassinoides* are identified in herbaria as *V. nudum*. This skews the results in the direction of "cold temperate," which is how Lens et al. (2016) scored *V. nudum*. Our recent analyses clearly distinguish these species, and show that *V. nudum* occupies only warm temperate forests (e.g., tupelo-cypress swamps) along the coastal plain of the Southeastern US, while *V. cassinoides* occupies colder forests to the north (Spriggs et al., 2019).

Third, our primary analyses included the locations and inferred biome states of five fossils. Four of the five appear to have occupied warm temperate forests, and this, along with the placement of the fossils in the tree, favors the conclusion that *Viburnum* evolved initially in warm forests. Although our results are not dependent on the inclusion of this information (Figs. S3 and S4), it does (and, we would argue, should) significantly increase the support for an origin in warmer forests.

Finally, we note that the Lens et al. (2016) conclusion that *Viburnum* lived ancestrally in cold climates is equivocal based on their analyses. With their preferred tree and scorings they concluded that "the ancestor of *Viburnum* has a PP of 0.52 to be of cold temperate origin (0.36 warm temperate origin and 0.12 tropical origin)." And, with *V. clemensiae* positioned as sister to the rest of *Viburnum*, it "has a PP of 0.45 to be of cold temperate origin (0.35 warm temperate and 0.21 tropical origin)". In contrast, our combined evidence analyses very strongly favor the hypothesis that *Viburnum* evolved in warm forests (Fig. 5).

*References*


Clement, W., M. Arikake, P. Sweeney, E. J. Edwards, and M. J. Donoghue. 2014. A chloroplast tree for *Viburnum* (Adoxaceae) and its implications for phylogenetic classification and character evolution. Amer. J. Bot. 101: 1029-1049.

Clement W. L. and M. J. Donoghue. 2011. Dissolution of *Viburnum* section *Megalotinus* (Adoxaceae) of Southeast Asia and its implications for morphological evolution and biogeography. Int. J. Plant Sci. 172: 559-573.

Eaton, D. A. R., E. L. Spriggs, B. Park, and M. J. Donoghue. 2017. Misconceptions on missing data in RAD-seq phylogenetics with a deep-scale example from flowering plants. Syst. Biol. 66: 399-412.

Egolf, D. R. 1962. A cytological study of the genus *Viburnum*. Journal of the Arnold Arboretum 43: 132-172.

Lens, F., R. A. Vos, G. Charrier, T. van der Niet, V. Merckx, P. Bass, J. A. Gutierrez, B. Jacobs, L. C. Doria, E. Smets, S. Delzon, and S. B. Janssens. 2016. Scalariform-to-simple transition in vessel perforation plates triggered by differences in climate during the evolution of Adoxaceae. Annals of Botany 118: 1043-1056.

Spriggs, E. L., W. L. Clement, P. W. Sweeney, S. Madriñán, E. J. Edwards, and M. J. Donoghue. 2015. Temperate radiations and dying embers of a tropical past: Evidence from *Viburnum* diversification. New Phytologist 207: 340-354.

Spriggs, E. L., D. A. R. Eaton, P. W. Sweeney, C. Schlutius, E. J. Edwards, and M. J. Donoghue. 2019. Restriction-site-associated DNA sequencing reveals a cryptic *Viburnum* species on the North American Coastal Plain. Syst. Biol. 68: 187-203.

Winkworth, R.C., and M. J. Donoghue. 2005. *Viburnum* phylogeny based on combined molecular data: implications for taxonomy and biogeography. Amer. Jour. Bot. 92: 653-666.

### Supplement 7. Supplementary Figures

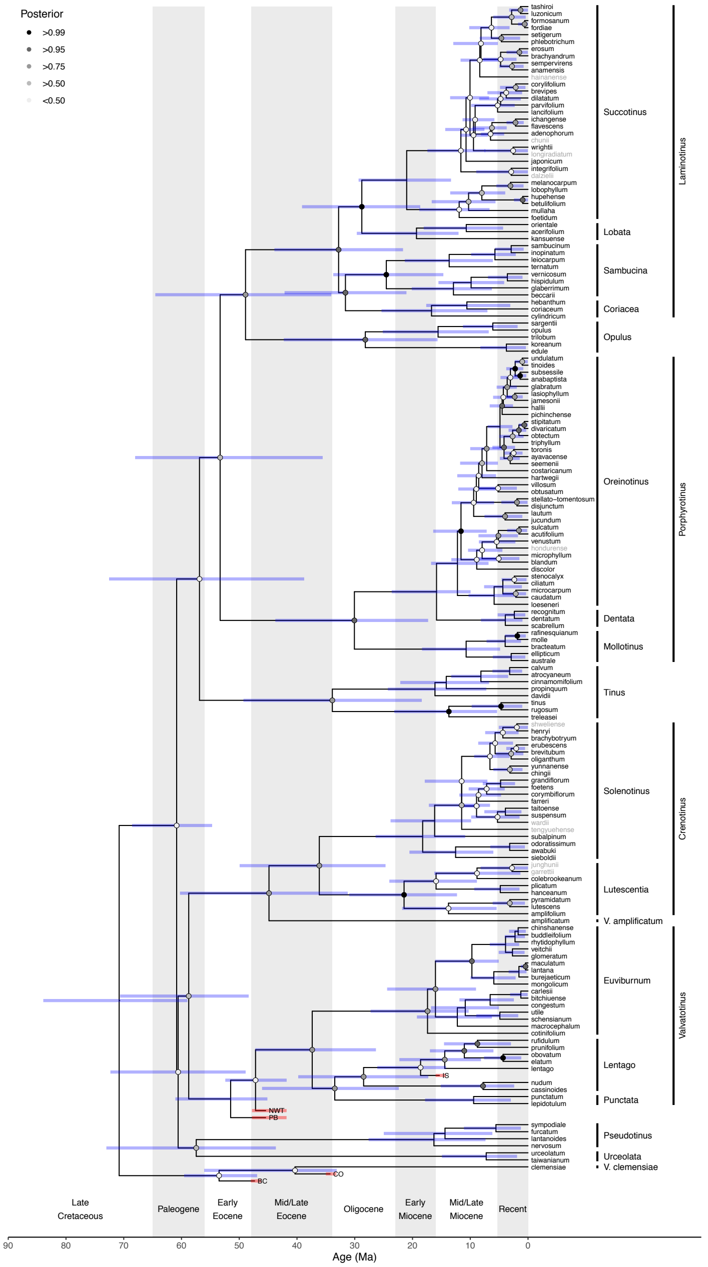


Figure S1. *Viburnum* divergence time estimates from the *Masked* dataset. Maximum clade credibility topology includes all 168 taxa. Node shading indicates posterior clade support. Node bars indicate 95% highest posterior density age estimates. Taxon labels for unsequenced extant taxa are colored gray.

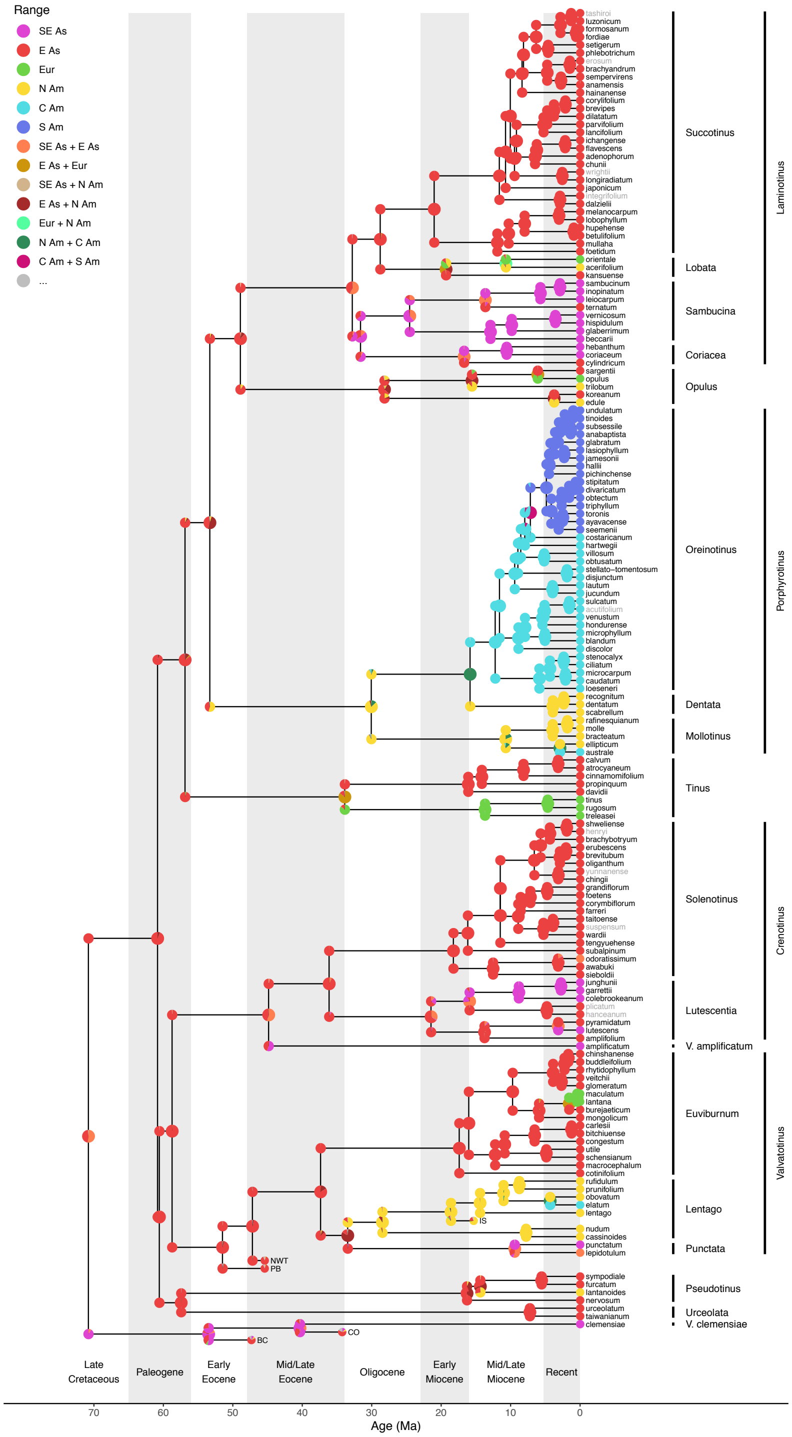


Figure S2. *Viburnum* ancestral range estimates from the *Masked* dataset. Node colors correspond to range states (legend). Range probabilities for the ancestral range and the daughter ranges at each internal node with pie charts. Taxon labels for unsequenced extant taxa are colored gray.

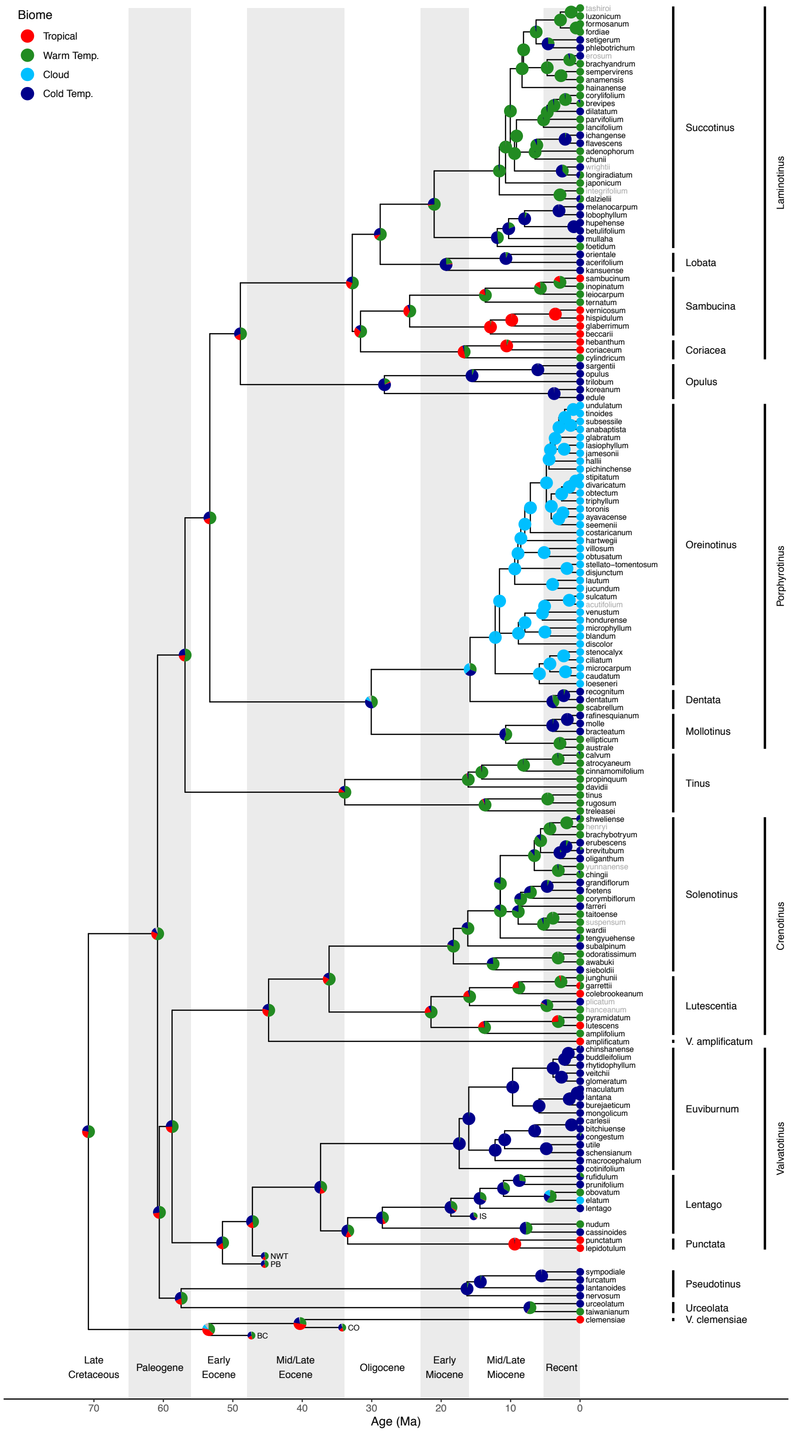


Figure S3. *Viburnum* ancestral biome estimates from the *Masked* dataset. Node colors correspond to forest biome affinity states (legend). Probabilities are given by pie charts at nodes. Taxon labels for unsequenced extant taxa are colored gray.

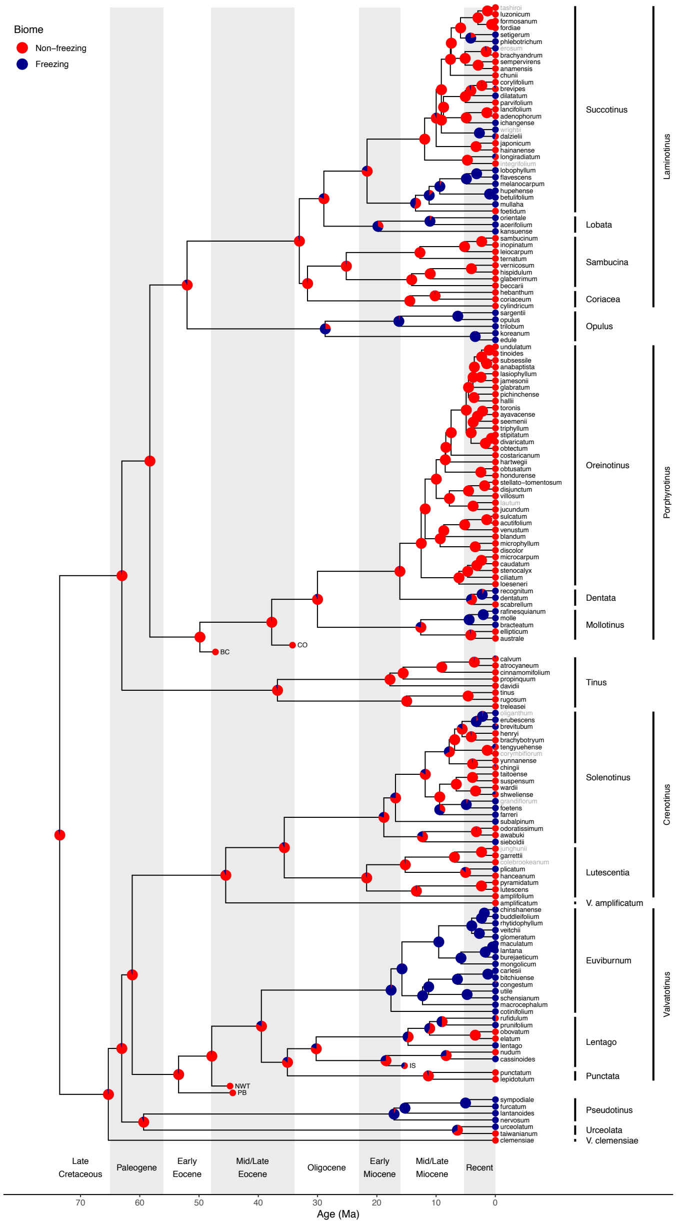


Figure S4. *Viburnum* ancestral biome estimates from the *Complete* dataset with two-state biome coding. Non-freezing biomes include tropical/subtropical (T), warm temperate/lucidophyllous (W), and cloud forest (Cl) biomes, while the freezing biome is the cold temperate (Co) biome. Node colors correspond to forest biome affinity states (legend). Probabilities are given by pie charts at nodes. Taxon labels for unsequenced extant taxa are colored gray.

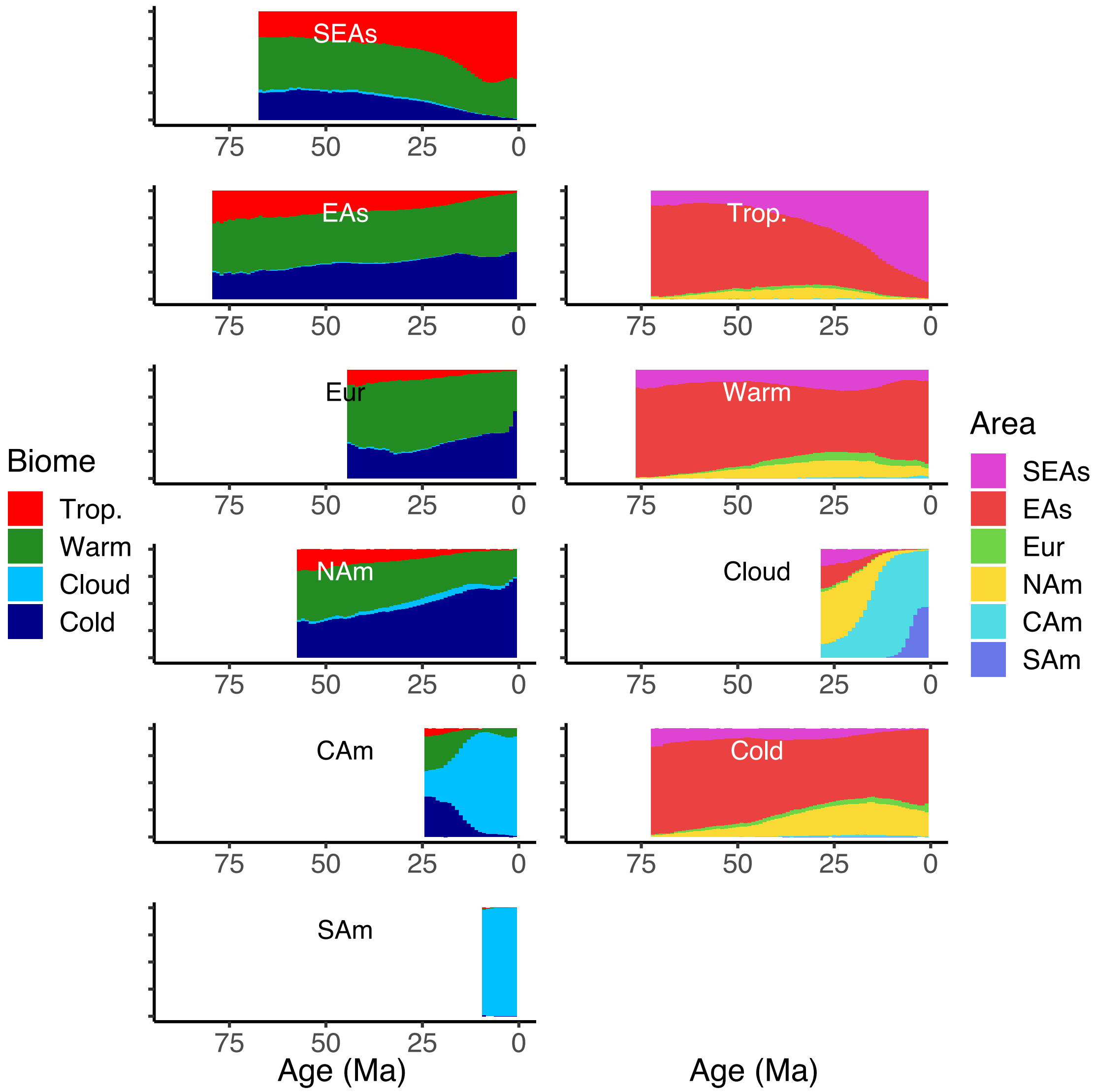


Figure S5. *Viburnum* biome and biogeography state frequencies through time as estimated from the *Masked* dataset. Subplots in the left column report the frequency of lineages across biome states given for all lineages with sampled descendants found within a particular region. Subplots in the right column are similar, except they report regional frequencies across lineages given a particular biome state. Time bins with too few posterior samples to guarantee accurate frequency estimates within ± 0.05 were marked as empty (see main text).

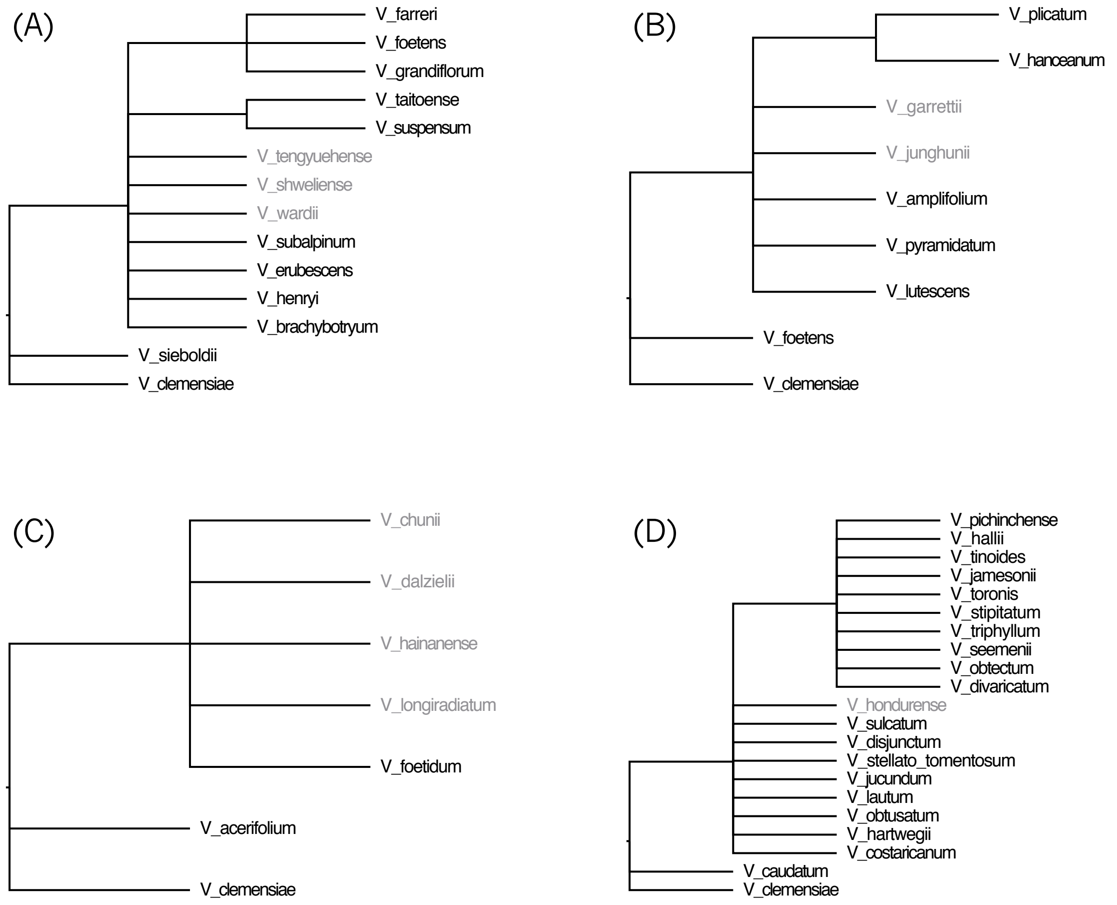


Figure S6. Morphology-based phylogenetic constraints for unsequenced species. Sequenced (black) and unsequenced (gray) species must obey the relationships defined by each phylogenetic constraint. Constraints are defined for: (A) *V. tengyuehense*, *V. shweilense*, and *V. wardii* in the *Solenotinus* clade; (B) *V. garretii* and *V. junghunii* in the *Lutescentia* clade; (C) *V. chunii*, *V. dalzielii*, *V. hainanense*, and *V. longiradiatum* in the *Succotinus* clade; and (D) *V. hondurense* in the *Oreinotinus* clade.

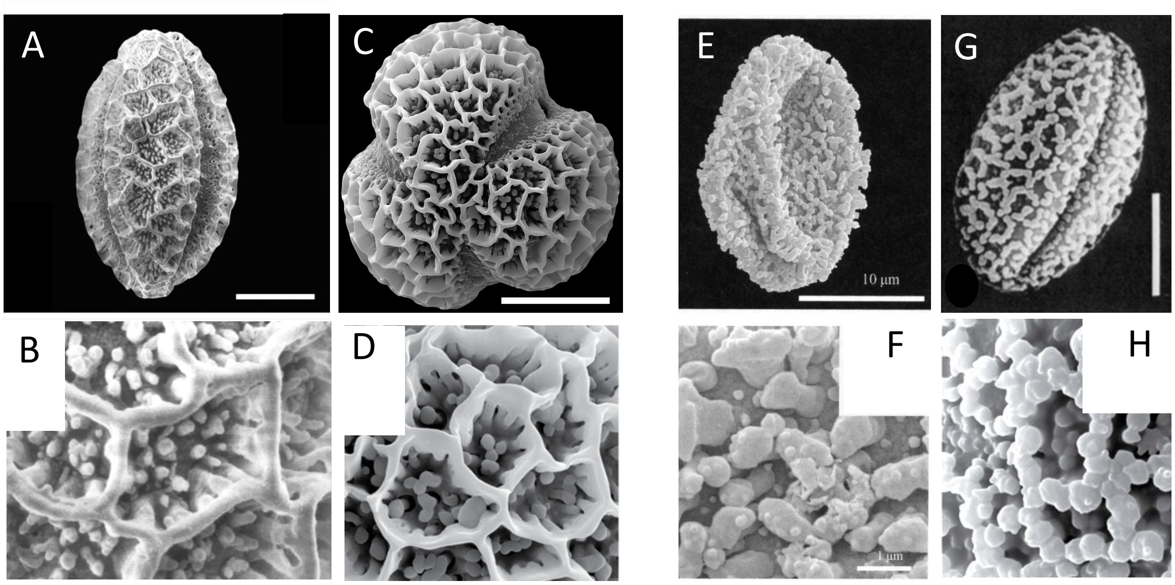


Figure S7. Scanning electron micrography of *Viburnum* pollen grains and exine morphology. A/B. Fossil pollen from British Columbia, ~48 Ma, exine type 1a (Manchester et al., 2015). C/D Modern *V. clemensiae* pollen, exine type 1a (Clement et al., 2014). E/F Fossil pollen from Iceland ~15 Ma, exine type 1c (Denk et al., 2011). G/H Modern *V. prunifolium* pollen, exine type 1c (Donoghue, 1985).

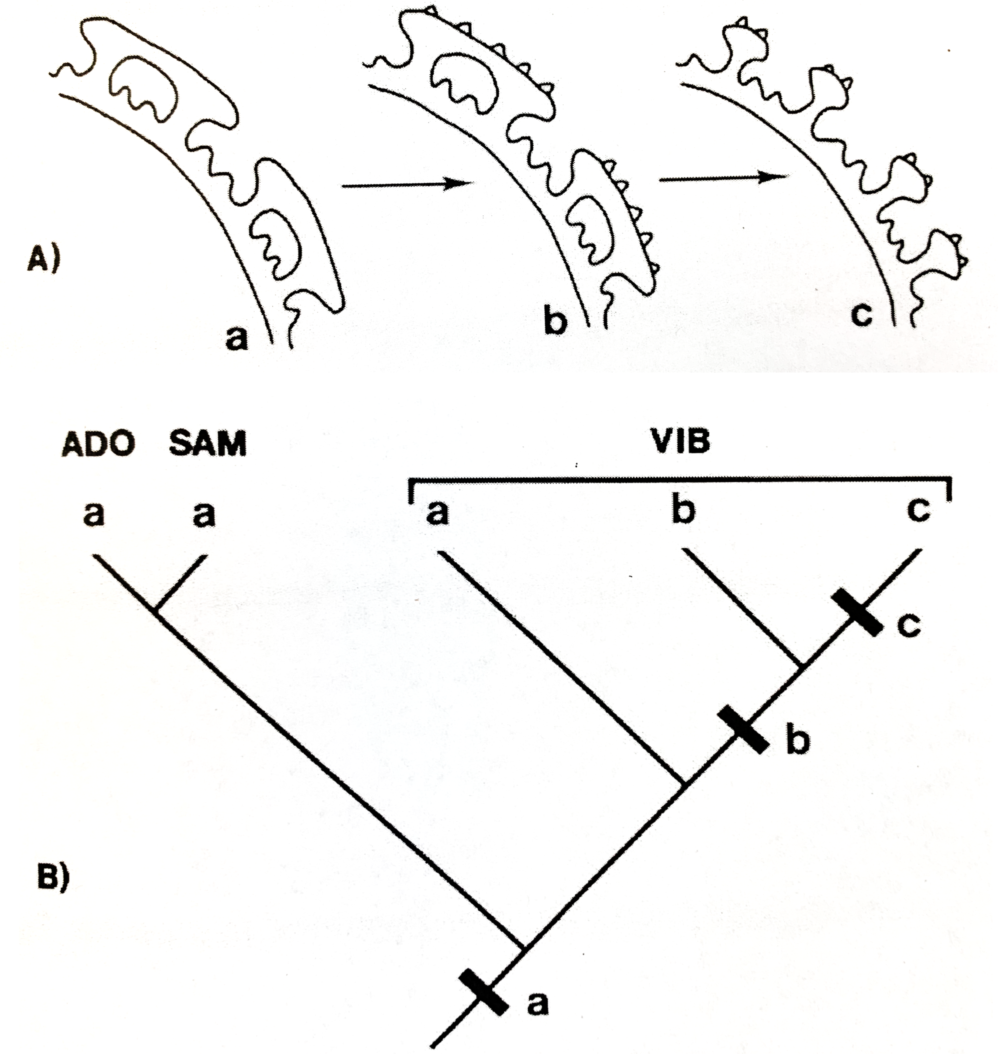


Figure S8. (A) Diagrams of cross sections through the three different pollen exine conditions found in *Viburnum*. In type Ia, there is a regular reticulum and the muri ("ridges") are smooth. In types Ib and Ic, the muri are scabrate ("bumpy"). In Ib there is a regular reticulum, whereas in Ic the reticulum is irregular and appears to be broken down. (B) As indicated by the arrows in (A), it is hypothesized that Ia is ancestral in *Viburnum* (and Adoxaceae more broadly). Type Ib is thought to have evolved from Ia, and type Ic from Ib. This has been confirmed in multiple phylogenetic analyses (e.g., Clement et al., 2014). ADO=*Adoxa*; SAM=*Sambucus*; VIB=*Viburnum*. Adapted from Donoghue (1985).
